# The long-range gene regulatory landscape of cerebellar granule neuron progenitors

**DOI:** 10.1101/2024.08.22.609103

**Authors:** Kimberley L. H. Riegman, Charlotte George, Danielle E. Whittaker, Mohi U. Ahmed, Haiyang Yun, Brian J. P. Huntly, David Sims, Cameron S. Osborne, M. Albert Basson

## Abstract

Neuronal specification, expansion and differentiation are tightly regulated by the concerted actions of transcription and chromatin modifying factors that are recruited to regulatory elements in the genome. Tissue-specific distal regulatory elements are typically located tens to hundreds of kilobases from the gene they regulate. Thus, to identify the distal enhancers that directly regulate a gene, information on the localisation of enhancers relative to the gene promoter in the nucleus is crucial. Cerebellar granule cell progenitors (GCps) are important transit amplifying neuronal progenitors, giving rise to the most abundant neuronal cell type in the brain. Many of the key factors that regulate fundamental developmental processes in GCps have been identified. For instance, the proneural transcription factor Atoh1 is essential for GCp specification, proliferation and differentiation and the ATP-dependent chromatin remodeller CHD7 is necessary for normal GCp proliferation and differentiation. However, both these factors are recruited to distal regulatory elements and the direct regulatory relationships between these factors, the enhancers they are recruited to, and the genes they regulate in GCps remain uncharacterised. To identify active, long-range gene regulatory interactions in GCps, we used promoter capture Hi-C (pcHi-C), and integrated pcHi-C data with ATAC-seq and ChIP-seq data. We present a rich dataset consisting of 46,428 interactions between 22,797 putative distal regulatory regions and 12,905 protein coding gene promoters in primary mouse GCps. Using VISTA-designated hindbrain enhancers as an example, we identify the genes most likely regulated directly by these enhancers and update their annotation accordingly. Motif enrichment analyses identified a significant enrichment of proneural transcription factor motifs in CHD7-regulated enhancers. Further analyses revealed co-localisation of Atoh1 and CHD7 at gene enhancers, suggesting a novel regulatory relationship between Atoh1 and CHD7 in controlling the expression of key genes in the GCp lineage. We used our data to identify >1,500 Atoh-regulated enhancers, contacting the promoters of 577 genes in GCps, and 197 enhancers of 22 genes that appear to be co-regulated by Atoh1 and CHD7. Co-immunoprecipitation experiments showed that Atoh1 and CHD7 proteins interact with each other. These findings support the emerging picture of CHD7 as an important gene regulatory co-factor for lineage-specific transcription factors. The pcHi-C data is presented as a useful resource to the community for investigating the function of long-range enhancers in the cerebellar GCp lineage.

## Introduction

Cellular differentiation is directed by the action of cell- and tissue-specific transcription factors. These factors are recruited to enhancer elements, which establish and maintain appropriate levels of gene expression at the correct time and place during development (Bulger & Groudine, 2010). Enhancers may be located at long distances from the genes they regulate and are brought in close physical proximity to these genes by means of chromatin looping (Bouwman & de Laat, 2015; Kadauke & Blobel, 2009; Schoenfelder & Fraser, 2019). Although a combination of local genomic features e.g. mono-methylation of lysine 4 on histone 3 (H3K4me1), acetylation of lysine 27 on histone 3 (H3K27ac) and accessible chromatin can identify active enhancer elements in the genome of different cell types (Calo & Wysocka, 2013), this information cannot identify the genes regulated by these enhancers. Chromatin conformation capture technologies like Hi-C (Lieberman-Aiden *et al*, 2009), and in particular an extension of Hi-C, promoter capture Hi-C (pcHi-C) that enriches for promoter interactions, overcome this limitation by identifying genomic regions that are in close spatial proximity to gene promoters (Mifsud *et al*, 2015; Schoenfelder *et al*, 2015). The frequency with which gene promotors and distal genomic regions are ligtaed in Hi-C is used as a measure to infer the likelihood of physical interaction in the nucleus and regulatory potential (Dekker *et al*, 2002). We used this approach to identify putative gene promoter-enhancer interactions in the developing cerebellum.

The cerebellum is essential for fine motor coordination and motor learning, and cerebellar hypoplasia or damage often leads to movement disorders like ataxia (Basson & Wingate, 2013). The cerebellum forms circuits with cortical regions that regulate movement, but also with regions involved in language and social processing (D’Mello & Stoodley, 2015), and may therefore have important functions in cognitive, affective and sensory processing (Allen & Courchesne, 2003). Cerebellar defects are associated with neurodevelopmental disorders, including Autism Spectrum Disorders (ASDs) (Fatemi *et al*, 2012; Wang *et al*, 2014), possibly by disrupting connectivity between the cerebellum and the medial prefrontal cortex (Kelly *et al*, 2020).

The most numerous neuronal cell type of the entire brain, cerebellar granule cells (GCs), are generated in the external germinal layer (EGL), a transient cell layer on the outer (pial) surface of the developing cerebellum, where transient amplifying divisions exponentially expand GCps (Consalez *et al*, 2020). In the mouse, GCps that form the EGL are specified in the rhombic lip from E13.5 (Ben-Arie *et al*, 1997; Machold & Fishell, 2005) and a clear EGL can be observed in the cerebellar anlage from E14 (Ben-Arie *et al*., 1997) of development. The GCp pool is amplified by proliferation from this stage and differentiation, induced by neurogenic factors like NeuroD1, can be detected from P0 onwards (Miyata *et al*, 1999).

After several cell divisions, GCps exit the cell cycle and initiate their differentiation programme. These post-mitotic progenitors migrate into the cerebellar cortex to form the internal granule cell layer (IGL), where these cells mature into GCs. Several transcription factors and chromatin regulators that control key steps in GCp differentiation have been identified. These factors are recruited to distal regulatory elements, although it is often unclear which genes they regulate. Glutamatergic cell specification in the cerebellum, as well as GCp expansion and differentiation are dependent on the Atoh1 pro-neural transcription factor (Klisch *et al*, 2011; Yamada *et al*, 2014). Atoh1 is preferentially recruited to distal enhancers, marked by H3K4me1 (Klisch *et al*., 2011). Thus, to identify genes directly regulated by specific Atoh1-bound enhancers, chromatin conformation capture approaches are required. One such Atoh1-regulated gene is *Zic4* (Zinc finger protein of cerebellum 4) (Klisch *et al*., 2011), which functions together with *Zic1* to maintain the progenitor state of GCps and therefore their proliferative capacity (Aruga *et al*, 1998; Blank *et al*, 2011; Grinberg *et al*, 2004).

In addition to transcription factors, ATP-dependent chromatin remodelling factors like CHD7 (Chromodomain Helicase DNA-binding factor 7) are recruited to distal enhancers, where they remodel chromatin structure to regulate gene expression (Schnetz *et al*, 2010). CHD7 is essential for normal expansion of GCps in the postnatal mouse cerebellum (Whittaker *et al*, 2017b) and deletion of *Chd7* from GCps results in striking cerebellar hypoplasia and polymicrogyria (Feng *et al*, 2017; Reddy *et al*, 2021; Whittaker *et al*., 2017b). *CHD7* haploinsufficiency is also sufficient to cause cerebellar hypoplasia and foliation defects both in mouse models and in the context of CHARGE syndrome in humans (Whittaker *et al*, 2017a; Yu *et al*, 2013).Thus, identifying CHD7- and Atoh1-regulated enhancers and their cognate genes is key to understanding fundamental transcriptional processes that control cerebellar GCp development.

The aim of this study was to identify active, long-range regulatory elements in cerebellar GCps and to use pcHi-C to identify genes directly regulated by these enhancers. To validate this dataset and provide further biological insights into gene regulatory mechanisms in GCps, we integrated our data with CHD7 ChIP-seq, differential chromatin accessibility and gene expression changes in *Chd7*-deficient GCps, as well as Atoh1 ChIP-seq and RNA-seq data from *Atoh1*-deficient GCps to identify genes directly regulated by Atoh1 and CHD7.

We report putative distal regulatory elements for >12,000 genes, identify CHD7- and Atoh1-regulated enhancer elements and show that these factors interact and likely co-regulate the expression of key genes in the GCp lineage.

## Results

### Promoter Capture Hi-C identifies long-range regulatory elements in cerebellar granule cell progenitors

To generate a genome-wide atlas of long-range promoter interactions at restriction fragment resolution in GCps, we generated two HindIII digested Hi-C libraries using freshly isolated proliferating mouse GCps. Promoter-containing fragments were captured from these Hi-C libraries with 39,684 biotinylated RNA baits (Yun *et al*, 2021a) which targeted 22,449 unique HindIII digested regions in the mouse mm10 genome (**Suppl. Table 1**). Of these targeted ‘bait’ fragments, 21,643 contained transcriptional start sites (TSSs), with 17,335 of these regions encompassing protein coding promoters. The pcHi-C libraries were processed using the Hi-C User Pipeline (HiCUP) (Wingett *et al*, 2015) to yield **∼**155 million and ∼134 million unique valid reads pairs for biological replicate one and biological replicate two, respectively. As is common in pcHi-C analysis (Freire-Pritchett *et al*, 2017), biological replicates were then bioinformatically combined into a ‘superset’ to increase sampling coverage, and significant interacting regions were identified using the Capture Hi-C analysis of Genomic Organisation (CHiCAGO) pipeline (Cairns *et al*, 2016), which identified 164,387 unique and significant interactions between a captured HindIII bait fragment and a distal genomic region. From hereon we will refer to these regions as promoter interacting fragments or PIFs. Of these interactions, 164,162 occurred in *cis* (intrachromosomal) and 195 in *trans* (interchromosomal). As we were interested in identifying unannotated *cis*-regulatory elements controlling transcription of genes important for cerebellar development, all trans-, bait-to-bait and non-protein-coding transcriptional start site (TSS) bait-to-PIF interactions were filtered from the dataset, resulting in 106,589 significant interactions for further analysis (**Fig. 1A**.**, Suppl. Table 2**). The median number of PIFs per bait was 5 and median distance of *cis* PIFs was ∼215kb, with 90% of interactions falling within ∼671kb (**Fig. 1B**), demonstrating the ability of this approach to identify long-range *cis*-regulatory interactions. These findings are in line with previously published pcHi-C datasets generated from other cell types (Mifsud *et al*., 2015; Siersbaek *et al*, 2017). To identify putative enhancer elements, we classified PIFs with respect to the presence of H3K4me1 and H3K27ac. This analysis revealed a 2.14-fold enrichment of H3K4me1 (**Fig. 1C)** and 2.69-fold enrichment of H3K27ac (**Fig. 1D)** at PIFs compared to what would be expected at random. Furthermore, heterochromatin regions, as defined by Ensembl regulatory annotations for brain (E14.5), were depleted from PIFs (**Fig. 1E**) and PIFs were highly enriched (4.52-fold) for active cerebellar enhancers identified by FANTOM5 (Dalby *et al*, 2018) (**Fig. 1F**). As an example, we identified three *Zic1* and *Zic4* PIFs, located ∼672kb (PIF1), ∼241kb (PIF2) and ∼592kb (PIF3) from the *Zic1* and *Zic4* promoters, which are located on the same restriction fragment in a head-to-head configuration (**Fig. 1G**). These regions were characterised by accessible chromatin (ATAC-seq) and enriched for H3K4me1 and H3K27ac histone modifications, consistent with these PIFs containing one or more active enhancer elements (**Fig. 1G)**. The *Zic1/Zic4* enhancer located ∼672kb upstream at chr9:90,686,000-90,708,000 (PIF1 in Fig. 1G) corresponds to the hs654 VISTA embryonic midbrain enhancer, indicating that this enhancer functions across different tissues and developmental time point to regulate *Zic1/Zic4* gene expression (Chen *et al*, 2024). The full set of PIFs with accessible chromatin (ATAC-seq), H3K4me1 and H3K27ac histone modifications are presented in **Suppl. Table 3**.

**Figure 1.**
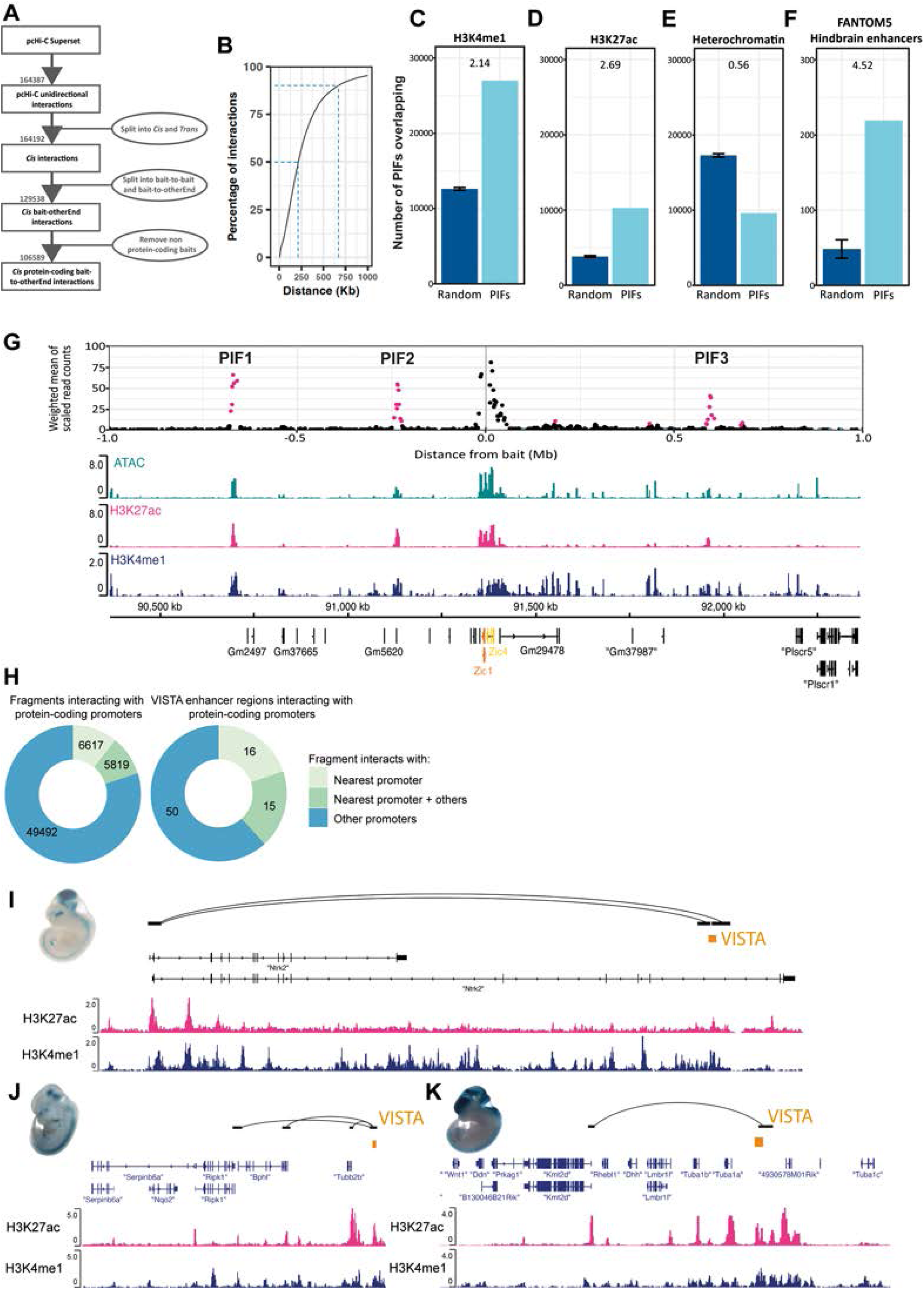
Promoter Capture Hi-C identifies long-range regulatory elements in cerebellar granule cell progenitors (GCps). **(A)** Schematic of the number of significant interactions (in grey) with a CHiCAGO score >=5 identified in our pcHi-C data-sets following each data filtering step (in ovals). Two independent replicates of pcHi-C library were pooled to create a ‘Superset’. Following filtering, 106589 *cis* protein-coding bait-to-otherEnd interactions (referred to as promoter interacting fragments or PIFs) were taken further for downstream analyses. **(B)** Graph demonstrating the cumulative percentage of significant interactions detected as genomic distance from the promoter capture baits increased. The first blue dotted line indicates the median interaction distance of *cis* PIFs (∼215kb) and the second blue dotted line indicates that 90% of interactions fell within 6711kb. **(C-F)** Bar charts illustrating the overlap of PIFs with (**C)** H3K4me1 ChIP-seq peaks, **(D)** H3K27ac ChIP-seq peaks, **(E)** heterochromatin regions defined by ensembl regulatory annotations for brain (E14.5) and **(F)** Fantom5 enhancers active in the cerebellum during p6-p9. Numbers above each graph indicate the enrichment (fold-change) of this overlap relative to 100 random subsets of *HindIII* fragments that have the same distribution in distances to the bait promoter regions as the significant PIFs. Error bars on random overlaps represent 95% confidence intervals from 100 permutations. **(G)** Top plot shows number of total normalised reads detected for each HindIII fragment interacting with the restriction fragment encompassing the *Zic1* locus from the pcHi-C dataset. Pink indicates locations PIFs identified as ‘significant’ with a CHiCAGO score >=5. Genome tracks show ATACseq (turquoise) peaks, and H3K27ac (pink) and H3K4me1 (navy) ChIPseq peaks for the same genomic region. **(H)** Plots illustrating the proportion of all PIFs (left) or PIFs containing VISTA hindbrain enhancers (right) that interact with either only the promoter of their nearest protein-coding gene (light green), those that interact with the promoter of their nearest protein-coding gene and interact with promoters of distal protein-coding genes (mid-green) and the proportion of PIFs interacting with one or more distal promoters that are not their nearest protein-coding gene (blue). Note that the majority of interactions identified are long-range and not with the nearest promoter. **(I-K)** Examples of interactions detected in our dataset for experimentally validated VISTA hindbrain enhancer regions showing **(I)** a VISTA enhancer that interacts with its nearest protein-coding gene promoter, *Ntrk2*, **(J)** a VISTA enhacer interacting with its nearest protein-coding gene promoter (*Tubb2*) as well as promoters for distal genes *Bphl* and *Ripk1* and **(K)** a VISTA enhancer that does not interact with its nearest protein-coding promoter but the distal promoter for *Kmt2d*. Insets show reporter activity for these VISTA enhancers in E11.5 or E12.5 mouse embryos (from VISTA enhancer site). Interacting HindIII fragments are shown in black with black arcs indicating fragments that localise together in the nucleus. Location of VISTA enhancer regions are shown in orange. Ensembl gene annotations are shown in dark navy, followed by H3K27ac (pink) and H3K4me1 (navy) ChIPseq data.

To confirm that PIFs identified by our analysis could function as enhancer elements in GCp cells, we cloned two fragments <u>(denoted #1.1 and #1.2,</u> Suppl. Fig. 1A), located in PIF1, the most significant *Zic1/Zic4* PIF according to CHiCAGO analysis Suppl. Fig. 1B), into the minimal promoter pGL3 luciferase vector (**Suppl Fig. 1E**). The VISTA enhancer browser (Visel et al., 2007) indicated that hs1203 largely overlapped #1.1 and element hs654 partially mapped to both fragments #1.1 and #1.2 (**Suppl. Fig. 1B, C, D**). Luciferase assays were performed in SHH-NPD cells, a GCp-derived cell line generated by Fults laboratory (Jenkins et al. 2014). To generate this cell line, GCps were infected to constitutively produce the mitogen SHH and maintain proliferation. Compared to the empty vector control, both #1.1 and #1.2 significantly induced luciferase activity in these cells (**Suppl Fig. 1F**) as expected for elements with enhancer activity in GCps.

When chromatin conformation or 3D organisation data is not available, studies typically assign regulatory elements to the nearest gene promoter. To test the validity of this approach, we compared this approach to the use of pcHi-C data to determine promoter-enhancer interactions. The majority (80%) of the PIF interactions we identified, bypassed the nearest gene promoter to interact with a more distal gene promoter. An additional 9% interacted both with the nearest gene as well as distal genes, with only 11% interacting exclusively with the nearest gene (**Fig. 1H**). To further demonstrate the power of this dataset to accurately assign *cis*-regulatory elements to their target gene, we investigated promoter interactions of experimentally validated hindbrain enhancers from the VISTA enhancer browser (Visel *et al*, 2007). At present, VISTA enhancers are annotated by their closest gene, with the assumption that these enhancers preferentially regulate the nearest gene. Of the 81 PIFs containing 65 experimentally validated VISTA hindbrain enhancers in our dataset, only 31 PIFs were found to ligate to promoter fragments from the nearest gene in our pcHi-C experiment. Of these, 16 were annotated in the VISTA database (see e.g. **Fig. 1I)**. We were able to assign additional distal interactions for 15 of these enhancers that demonstrated interactions with their nearest gene (see e.g. **Fig. 1J**) and we could assign distal enhancers for the remaining 50 PIFs containing VISTA enhancers that did not appear to interact with the closest gene in our data (see e.g. **Fig. 1K**). This suggests, as reported by several other studies (Chen *et al*., 2024; Li *et al*, 2012; Mifsud *et al*., 2015; Sanyal *et al*, 2012), that nearest-gene annotations do not accurately annotate the majority of enhancer elements. In our case, 65/81 (80%) of PIFs containing VISTA enhancers were found to have additional promoter interactions beyond their nearest annotated gene. The gene annotations for the 81 PIFs containing VISTA hindbrain enhancers are presented in **Suppl. Table 4.** In summary, >80% of enhancers appeared to regulate distal genes in GCps, although we cannot rule out that some additional interactions with the nearest gene promoters might be identified in our pcHi-C dataset with deeper sequencing or alternative interaction calling algorithms that prioritise proximal interactions.

### Integration of pcHi-C data with ATAC-seq improves the resolution of PIFs containing distal regulatory elements

As PIFs are relatively large (median size 4.2 kb) and frequently found in clusters around open chromatin regions in our data we integrated DNA accessibility (ATAC-seq) data with pcHi-C to identify enhancers. ATAC-seq data from P7 GCps identified >112,000 500bp wide ATAC-seq peak summits across the GCp genome and intersection with our pcHi-C data demonstrated that ∼27% (30,296/112,341) of these fell within PIFs. A total of 37% (22,797/61,928; **Fig. 2A**) of PIFs and 44% of all PIF-promoter interactions (46428/112,341; **Fig. 2B**) contained an ATAC-seq peak summit. Many remaining PIFs appear to flank PIFs with ATAC-seq peaks. The median distance of a PIF without an ATAC-seq peak from a PIF with an ATAC-seq peak was 4.5 Kb (**Fig 2C**), which is just over the median length of a PIF in the dataset (4.2Kb). 66% of PIFs without ATAC-seq peaks were found within 5 restriction fragments of a PIF containing an ATAC-seq peak (**Fig. 2D**). Together, these observations suggest that the majority of PIFs that do not contain an ATAC-seq peak are proximal to PIFs with accessible genomic regions and that focusing on those regions with accessible sites will enable a more accurate identification of regulatory sites than using PIFs identified by pcHi-C alone.

**Figure 2.**
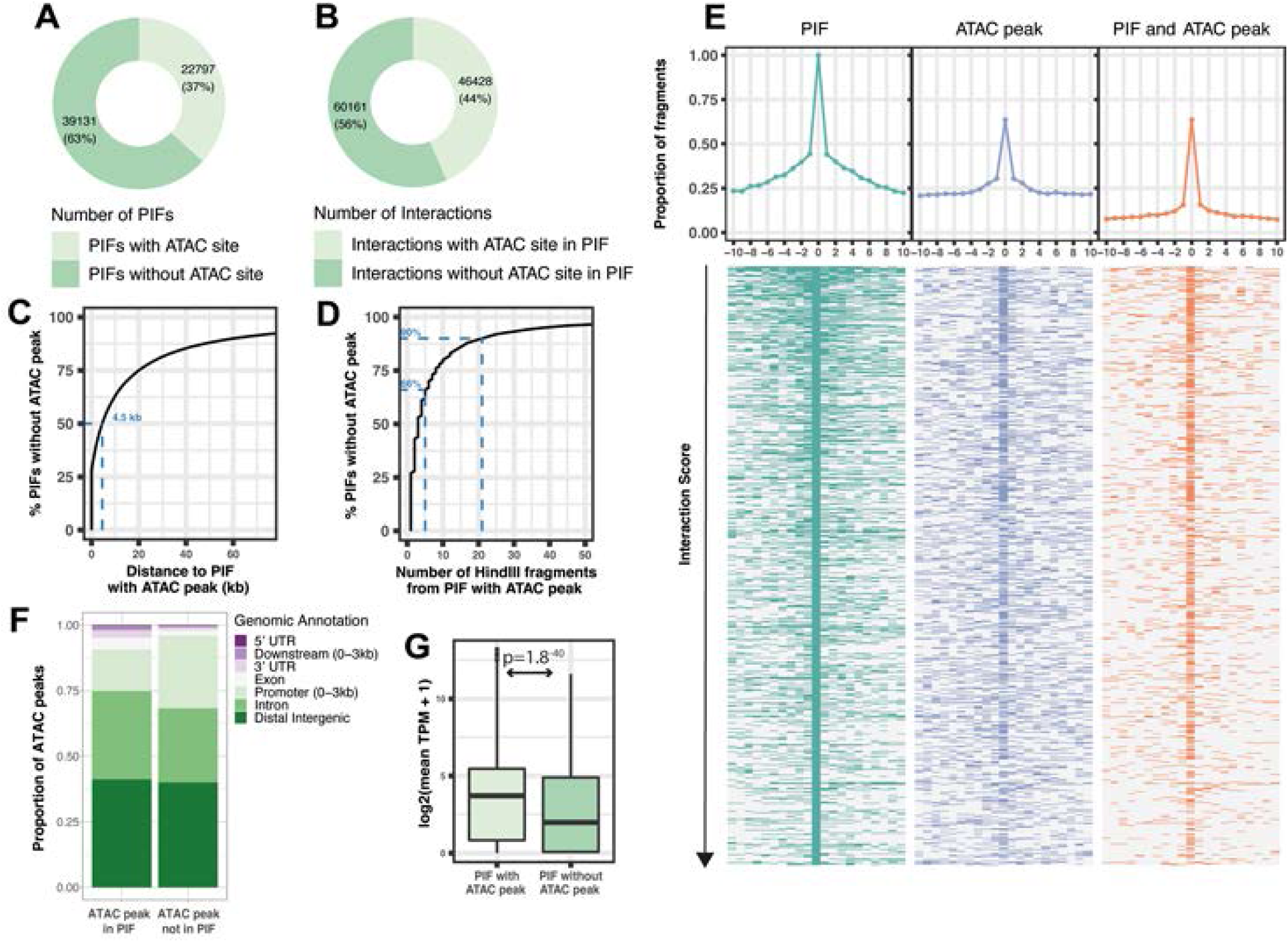
The majority of promoter interacting fragments (PIFs) are within 5kb of a PIF that contains an accessible ATAC peak. Plots illustrating the number and proportions of **(A)** PIFs containing an accessible ATAC peak and **(B)** promoter-PIF interactions with a PIFs containing an accessible ATAC peak. Cumulative percentage graphs demonstrating the genomic distance **(C)** and number of HindIII restriction fragments **(D)** PIFs without accessible regions (without ATAC peak) are located from their nearest accessible PIF (PIF with ATAC peak). The blue dotted line in **(C)** indicates the median distance between a PIF without an ATAC peak and a PIF with an ATAC peak is 4.5 kb, whilst 66% of PIFs without an ATAC peak are within a range of 5 genomic HindIII fragments from an accessible PIF **(D)**. **(E)** PIF and accessibility status of the 10 upstream and downstream genomic HindIII restriction fragments surrounding the ‘lead PIF’ with the highest and most significant CHiCAGO score for each captured promoter. Central point of heatmaps and coverage plots show the lead PIF (fragment 0). Heatmaps show the PIF status (left, green in heatmap if genomic fragment is PIF, white if not a PIF), ATAC status (middle, purple in heatmap if genomic fragment contains ATAC peak, white if no ATAC peak) and combined PIF and ATAC status (right, orange if genomic fragment is identified as a PIF and contains a ATAC peak, while if not). Each row represents the primary PIF for a different captured promoter, with rows ordered by descending CHiCAGO score, i.e most significant PIF at the top. Coverage plots at the top of each heatmap indicate the overall proportion of fragments flanking the primary PIF on either side that are also designated as PIFs (left, green), containing ATAC peaks (purple, middle) or PIF with an ATAC peak within it (orange, right). **(F)** Genomic location of ATAC peaks within and not in PIFs. **(G)** Boxplot showing the distributions of the average expression levels (TPM) of genes with promoters in proximity to PIFs containing ATAC peaks compared to promoters with PIFs that do not contain accessible regions.

To further confirm the relationship between accessibility and PIFs for each promoter in the pcHiC dataset we identified the ‘lead PIF’ with the highest CHiCAGO score for each promoter region and plotted the PIF and chromatin accessibility status of the upstream and downstream flanking HindIII restriction fragments either side (**Fig. 2E**). The ‘lead PIF’ with the highest CHiCAGO score for each bait tended to be surrounded by significant PIFs on either side which decreased as the number of restriction fragments from the primary PIF increased (**Fig. 2E left panel**) and demonstrated a strong co-localisation with accessible ATAC-seq peaks **(Fig. 2E right panel**). We therefore focused our analysis on the ATAC-seq sites within PIFs rather than on the PIF fragments themselves as these sites are likely to contain the distal regulatory sequences driving promoter interactions **(Suppl. Table 5)**.

Annotation of the ATAC-seq peaks within PIFs revealed that the majority of these accessible sites were located in distal intergenic regions or intronic regions **(Fig. 2F)**, whilst ATAC-seq peaks that did not colocalise with our PIF dataset contained a slightly higher proportion of promoter regions as would be expected due to the removal of ‘bait-to-bait’ promoter-to-promoter interactions from our PIF dataset. Integration of this data with bulk RNA-seq data allowed the examination of the expression levels of 14,403 protein-coding genes that could be linked to PIFs. The 12,354 genes that were linked to a PIF containing an ATAC-seq peak were found to have a higher median expression level than the 2,049 genes that had PIFs that did not coincide with ATAC-seq peaks (**Fig. 2G**).

Gene Ontology analysis of genes with accessible PIFs revealed a significant enrichment for 119 biological processes with embryonic organ morphogenesis (GO:0048562; *P =* 1.42 x 10^-12^), axon guidance (GO:0007411; *P =* 1.24x10^-9^) and neuron projection guidance (GO:0097485; *P =* 1.24x10^-9^) in the top 5 most significant terms, whilst no significant processes were enriched in the genes without accessible PIFs (**Suppl. Table 6**). Our subsequent investigations therefore focused on PIFs containing accessible regions, and the ATAC-seq peaks within these PIFs to identify the sequences and transcription factors that might be driving promoter-enhancer interactions in GCps.

### The chromatin remodeller CHD7 regulates gene expression in GCps by promoting chromatin accessibility at distal enhancer elements

As the chromatin remodeller CHD7 is primarily recruited to distal gene enhancers, rather than promoters (Reddy *et al*., 2021; Schnetz *et al*., 2010), it has not been possible to identify the genes directly regulated by CHD7 in GCps. We therefore integrated our pcHi-C and ATAC-seq datasets with ChIP-seq data to identify CHD7-regulated genes in GCps.

Differential accessibility analysis of the 112,341 accessible chromatin regions in the GCp genome, identified 5,369 regions that showed significant changes (p-adj<0.05) in accessibility in *Chd7*-deficient GCps (**Fig. 3A**), the majority of which (4,663/5,369) displayed decreased accessibility when *Chd7* is depleted, as previously described (Whittaker *et al*., 2017b). Overlapping these data with the 30,296 ATAC-seq sites that were present within PIFs (**Fig. 3A**), we were able to assign 1,465 (27%) of the ATAC-seq regions affected by *Chd7* deficiency to a gene promoter using the pcHi-C dataset. Several of the CHD7-regulated elements were in contact with multiple gene promoters, such that 2,371 protein coding genes appeared to have distal sites with chromatin accessibly regulated by CHD7. As some of the accessibility changes at these putative enhancer elements may be indirect, we further integrated the ATAC-seq data with RNA-seq from *Chd7*-deficient GCps (Whittaker *et al*., 2017b) and CHD7 ChIP-seq data (Feng *et al*., 2017) to identify enhancers that are likely directly regulated by CHD7. This analysis identified 210 enhancers occupied by CHD7 that changed in accessibility on CHD7 depletion (**Fig. 3A**), with 66 of these interacting with the promoters of 63 genes that were differentially expressed in *Chd7*-deficient GCps (**Fig. 3B**). These genes represent direct CHD7 targets with high confidence. Over-representation analysis confirmed an enrichment of genes linked to nervous system development (Padj=9.3x10^-4^) and neurogenesis GO:biological processes (Padj=5.1x10^-3^) (**Suppl. Table 7**) with 19 of these 63 genes upregulated and 44 downregulated in *Chd7*-deficient GCps (**Fig. 3B, Suppl. Table 8**).

**Figure 3.**
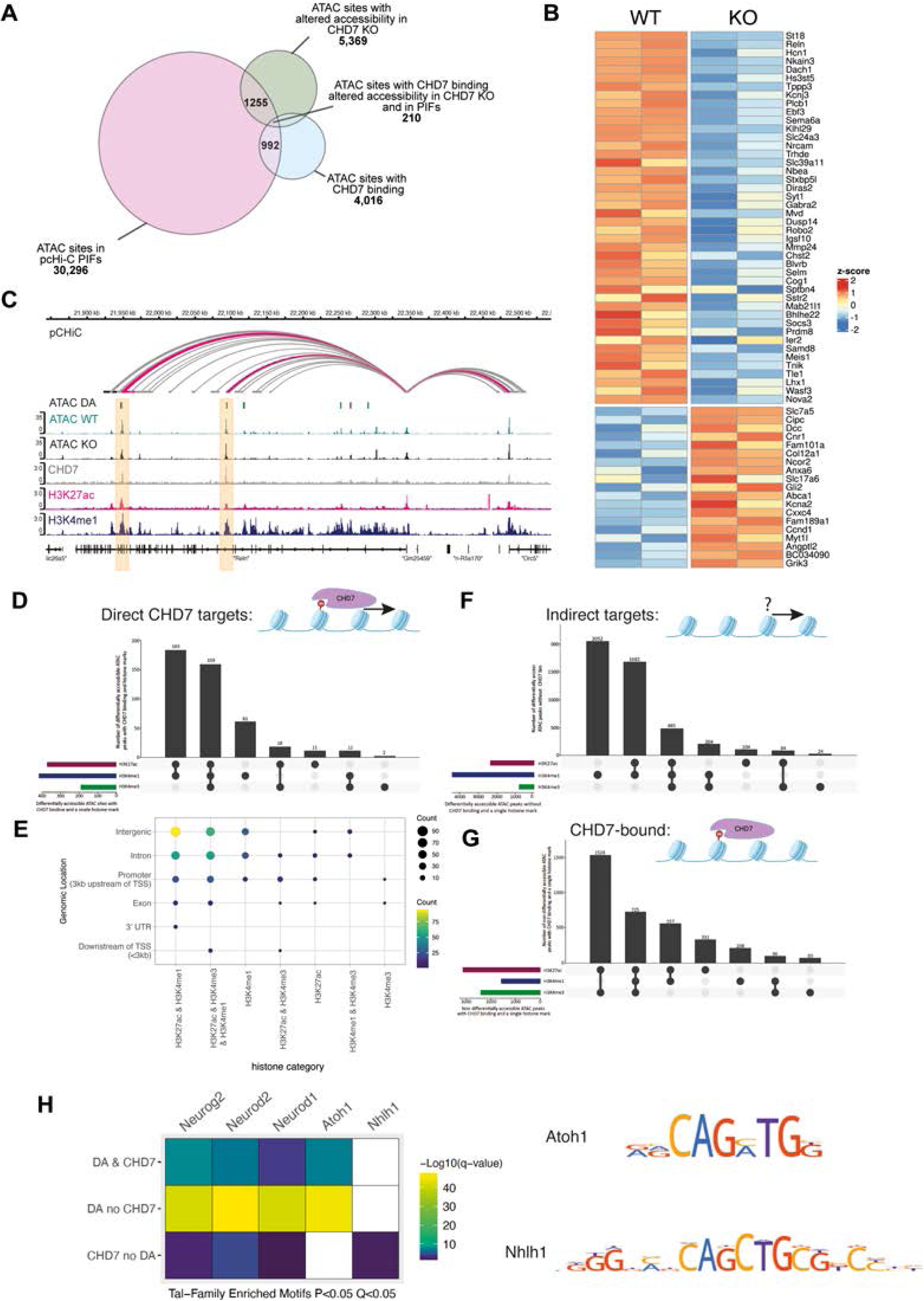
The chromatin remodeller CHD7 regulates gene expression in GCps by promoting chromatin accessibility at distal enhancer elements. **(A)**Venn diagram showing the number of ATAC accessible chromatin peaks found within PIFs (30,296 – pink), with altered accessibility in *Chd7*-deficient GCps (5369 – green), that overlap CHD7 ChiPseq peaks (4016 – blue) and the overlap of these subsets. 210 promoter-proximal ATAC peaks with CHD7 recruitment and which change accessibility in *Chd7*-deficient GCps are considered direct CHD7 targets. **(B)** Heatmap of direct CHD7 target genes which are differentially expressed in *Chd7*-deficient GCps and have at least one PIF which overlaps a CHD7 binding site that shows altered accessibility in *Chd7*-deficient GCps. Values depict z-score of normalized, log2 transformed and scaled RNAseq data between p7 WT (n=2) and *Chd7*-deficient KO (n=2) GCps. **(C)** Genome tracks at the *Reln* locus (chr5:21843156-22533049) showing the location of three putative *Reln* enhancers with CHD7 binding signal and decreased accessibility in *Chd7*-deficient GCps identified 395kb, 394kb and 249kb downstream of the *Reln* promoter (within the two orange shaded regions). Promoter proximal PIFs are designated by arcs (grey for any PIF with significant CHiCAGO score ≥5, pink for PIFs that also contain an ATAC peak that overlaps with CHD7 binding signal. Regions of differential chromatin accessibility (p-adj < 0.05) are displayed in the interval track (ATAC DA) showing 500bp regions of decreased (green) and increased (purple) accessibility in *Chd7*-deficient GCps between the WT ATAC (n=3) and *Chd7*-deficient (KO) ATAC signal (n=3). A representative ATAC signal track for WT (green) and CHD7 KO (black) GCps are shown above the ChIPseq signal tracks for CHD7 (grey), H3K27ac (pink) and H3K4me1 (dark blue). **(D)** UpSet plot showing the predominant combinations of chromatin modifications present at ATAC peaks that overlap CHD7 ChIPseq peaks and change accessibility in C*hd7*-deficient GCps. Number of ATAC peaks overlapping individual chromatin marks are shown in smaller horizontal bars on left of plot, vertical bars of main plot indicate number of ATAC peaks with combinations of H3K27ac, H3K4me1 and H3K4me3 marks as denoted in the panel below x-axis. Diagram illustrates CHD7-mediated chromatin remodelling (arrow). **(E)** Dot plot showing the number of ATAC peaks at different genomic locations (y-axis) for different combinations of chromatin marks (x-axis) from **(D)** for the direct CHD7 target sites that have ATAC peaks which change in accessibility in C*hd7*-deficient GCps and overlap CHD7 ChIPseq signal. **(F)** UpSet Plots illustrating predominant chromatin mark combinations for ATAC peaks that change in accessibility upon C*hd7*-depletion but do not exhibit CHD7 recruitment. Diagram illustrates chromatin remodelling (arrow) by an unknown factor (?). **(G)** aUpSet plots of ATAC peaks with CHD7 recruitment but no changes in accessibility upon C*hd7*-depletion. Diagram illustrates CHD7 presence without detectable chromatin remodelling. **(H)** Heatmap of significant SEA motif-enrichment results (p<0.05, q<0.05) for TAL-family transcription factors for genomic regions with ATAC peaks that are direct targets of CHD7 with CHD7 occupancy and change in accessibility in C*hd7*-deficient GCps (DA & CHD7), those that alter accessibility upon CHD7 depletion but do not display CHD7 occupancy in WT (DA no CHD7) and those which display CHD7 recruitment but do not alter accessibility upon CHD7 depletion (CHD7 no DA). White boxes denote no statistically significant enrichment and consensus motifs for TAL family members that show differential enrichment across the subsets are shown on the right. Also see Suppl. Fig. 1.

We were able to identify accessible distal regulatory elements for 632 (76%) of the 830 genes that are differentially expressed in the *Chd7*-deficient GCps. In addition to the 63 genes that were linked to distal sites both bound by CHD7 and demonstrating altered accessibility (direct CHD7 targets), we identified a further 206 genes with a) CHD7 binding at distal elements but no changes in accessibility of these regulatory regions (64 genes, putative direct CHD7 targets), b) chromatin accessibility changes with no CHD7 binding (99 genes) or c) with neither CHD7 recruitment or accessibility changes detected (43 genes), the latter two groups representing indirect CHD7 targets (**Suppl. Table 8**).

The cerebellar hypoplasia and foliation changes in CHD7 mutants have been linked with dysregulation of the *Reln* gene (Whittaker et al., 2017a). We therefore asked if our pcHi-C data can identify any CHD7-regulated *Reln* enhancers. We identified three putative intronic Reln enhancers: 396kb, 395kb and 249kb downstream of the *Reln* TSS which showed altered accessibility in *Chd7*-deficient GCps. The region 395kb downstream, located within intron 43 of the *Reln* gene, is also bound by CHD7 and therefore a CHD7-regulated *Reln* enhancer. Importantly, the putative *Reln* enhancer in intron 8 highlighted in our previous publication as a putative CHD7-regulated enhancer (Whittaker *et al*., 2017b), does not show any interaction with the *Reln* promoter in our pcHi-C data, suggesting that this is not a CHD7-regulated enhancer in GCps, although we cannot exclude that deeper sequencing will identify such interactions.

We further characterised the accessible genomic regions described in **Fig. 3A** as active enhancers (H3K27ac+, H3K4me1+), promoters (H3K27ac+, H3K4me3+), regulatory elements (H3K27ac+, H3K4me1+, H3K4me3+), or poised enhancers (H3K4me1+) (Creyghton *et al*, 2010; Gates *et al*, 2017). Genomic regions likely to be directly regulated by CHD7 (differential accessibility (DA) ATAC-seq sites and CHD7 binding sites) were marked as active enhancers (183/445=41%) and regulatory elements (159/445=36%), with 61/445 = 14% marked as “poised” enhancers by H3K4me1 alone (**Fig. 3D**). Active and poised enhancer elements were mostly found in intergenic and intronic regions **(Fig. 3E).** ATAC-seq sites with altered chromatin accessibility but no clear CHD7 binding, were predominantly classified as poised (2,052/4,635=44%) or active enhancers (1,682/4,635=36%) (**Fig. 3F**). ATAC-seq sites with CHD7 binding but no change in chromatin accessibility comprised mostly of active (H3K27ac) elements with H3K4me3 (2,251/3443=65%) (**Fig. 3G**), suggesting that the presence of CHD7 at active promoters was rarely associated with clear alterations in chromatin structure. These data are consistent with previous studies which demonstrated that CHD7 primarily regulates enhancers.

### CHD7-regulated enhancers are enriched for proneural transcription factor motifs

CHD7 has been shown to cooperate with tissue-specific transcription factors in different cell types (Engelen *et al*, 2011; Hsu *et al*, 2020; Stathopoulou *et al*, 2023). We reasoned that a comparison of transcription factor motifs enriched within CHD7-regulated enhancers might identify transcription factors that cooperate with CHD7 in GCps.

We used the MEME suite Simple Enrichment Analysis (SEA) tool (Bailey & Grant, 2021b) to identify motifs from the HOCOMOCO Mouse v11 CORE database that were overrepresented in ATAC-seq regions of interest in our PIFs, compared to all ATAC-seq sites in the cells. This analysis revealed a highly significant enrichment of E-box motifs corresponding to putative consensus sites for proneural bHLH TAL-family of proteins Neurog2, Neurod2, Neurod1 and Atoh1 in elements that show CHD7 recruitment as well as differential chromatin accessibility in CHD7-deficient GCps (**Fig. 3H**). The even stronger enrichment in regions without obvious CHD7 recruitment might indicate that some CHD7-regulated elements are being missed due to poor CHD7 ChIP-seq. All these proneural factors were expressed in GCps, and their expression was not altered in *Chd7*-deficient GCps, ruling out the possibility that these effects are mediated via changes in the expression of proneural transcription factors (**Suppl. Fig. 2)** . Together, these findings suggests that CHD7 preferentially regulates chromatin accessibility at enhancer regions containing these motifs and therefore may work in concert with proneural factors to regulate gene expression in cerebellar GCps. Intriguingly, the Atoh1 motif was specifically enriched at those elements that showed chromatin accessibility changes, whilst Neurod1, Neurod2 and Neurog2 motifs were also present in elements that showed no chromatin accessibility changes in *Chd7*-deficient GCps (**Fig. 3H**). These findings suggest that CHD7 activity in proliferating GCps is preferentially localised to Atoh1-regulated enhancer regions, whilst chromatin at Neurod1-regulated regions may not yet be remodelled by CHD7. This finding agrees with our finding that *Chd7*-deficiency preferentially affects GCp proliferation, the primary process regulated by Atoh1 and not differentiation, the process initiated by Neurod1 (Whittaker *et al*., 2017b). Regions with CHD7 recruited but without clear changes in chromatin accessibility in CHD7-deficient GCps were enriched for the Tal-Family member NHLH1 (**Fig. 3H**) along with other transcription factors including NF1A, bHLH ZIP factors (SREBF1/2, 3 Zn finger families (SP2-5, EGR1/2, ZIC1, ZBTB), E2F-related, AP2, TCF7-related and RFX-related factors (**Suppl. Fig. 2).**

Regions bound by CHD7 showed a lack of enrichment for TALE-type homeodomains, SMAD, FOS, JUN and PAS motifs that were enriched in PIFs with altered chromatin accessibility but no clear CHD7 ChIP-seq signals, suggesting that these transcription factors motifs were engaged indirectly in *Chd7*-deficient GCps (**Suppl. Fig. 2)**. We also noted that the genes encoding NFYB, NHLH1, SREBF2 and E2F1 were downregulated in *Chd7*-deficient cells whilst *Smad3* and *Mafb* gene expression was upregulated (**Suppl. Fig. 2)**, suggesting that some effects on the activity of these enhancers and altered expression of their cognate genes in *Chd7*-deficient GCps may be secondary to transcription factor expression changes, rather than a direct effect of CHD7 loss. Together, these finding strongly suggested that CHD7 may function as an important co-factor of the lineage-specific proneural transcription factor Atoh1 in GCps. Given the critical role for Atoh1 in the GCp lineage, we next sought to determine if CHD7 is recruited to Atoh1-regulated enhancers.

### Long-range enhancers bound by the key GCp lineage proneural factor Atoh1

Atoh1, a proneural, basic helix–loop–helix transcription factor, is essential for GCp fate specification and proliferation (Ben-Arie *et al*., 1997). Previous studies have suggested that Atoh1 functions at H3K4me1+ enhancers (Klisch *et al*., 2011), however, the genes directly regulated by these enhancers have not been identified. We reanalysed ChIP-seq and RNA-seq datasets from Klisch et al. and identified 7406 accessible regions bound by Atoh1 in 6890 PIFs proximal to promoters of 7090 protein-coding genes (**Fig. 4A**). Of these, 1,854 accessible ATAC-seq sites could be assigned to 577 genes that are differentially expressed (adjusted *P*-Value < 0.05) in *Atoh1*-deficient cells (**Fig. 4B**, **Suppl. Table 9**). Consistent with our analysis of our full data-set and VISTA enhancers reported above, only ∼14% of Atoh1 bound ATAC-seq sites (263/1854) interacted with only their nearest protein-coding gene promoters (**Fig. 4C)**.

**Figure 4.**
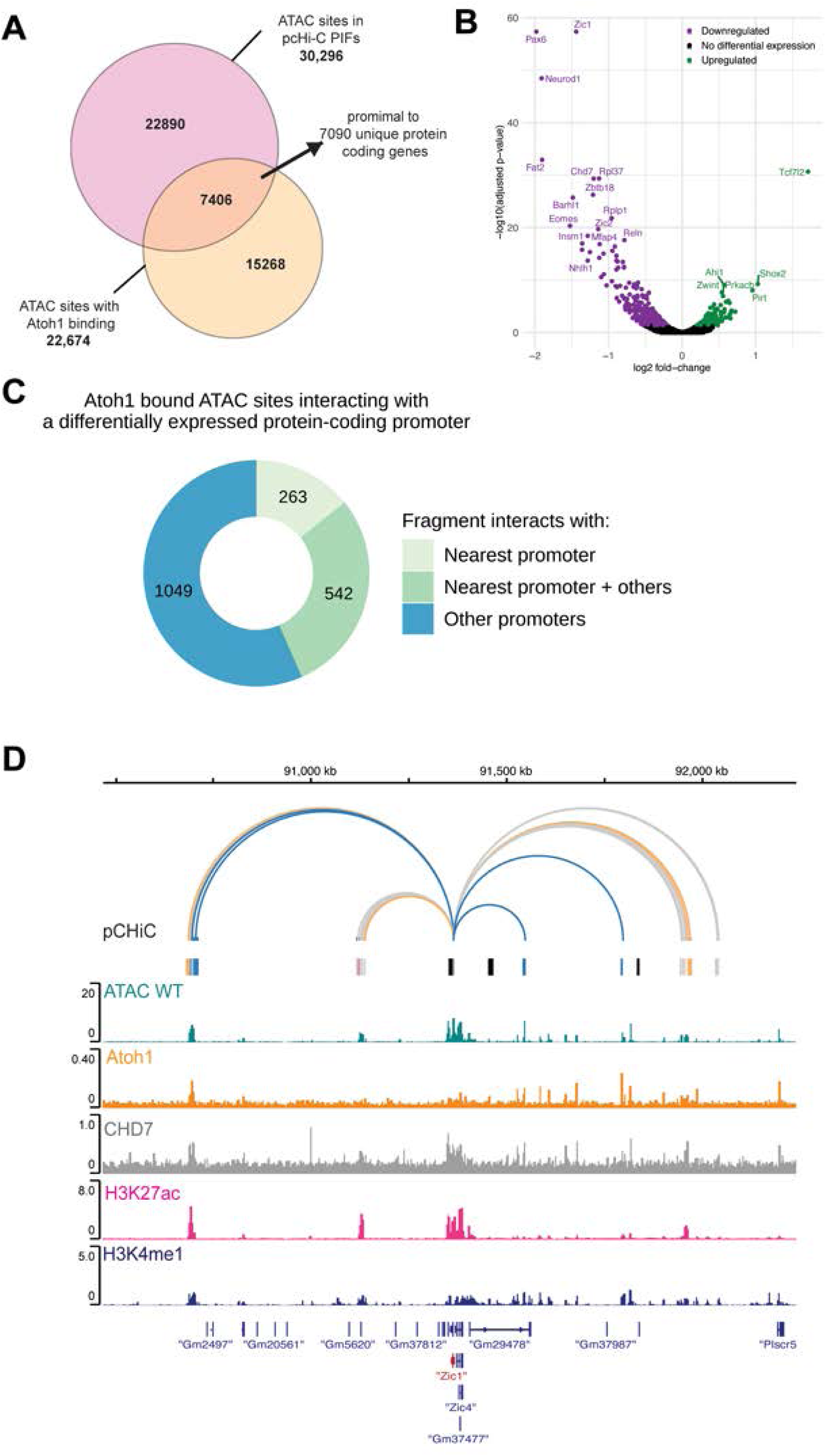
Atoh1-regulated distal regulatory elements and gene targets in primary GCps. (**A**) Schematic venn diagram of the number of accessible ATAC-seq sites in PIFs that overlap with accessible ATAC-seq sites that display Atoh1 occupancy in GCps. The 7406 accessible sites in PIFs that have Atoh1 occupancy are proximal to promoters of 7090 protein coding genes, 599 of which are differentially expressed in Atoh1-deficient GCps. (**B**) Volcano plot showing magnitude of multiple testing corrected p-value and log2-fold expression changes of 599 protein coding genes in the Atoh1-deficient cerebellum that have been linked to distal regulatory regions with accessible sites that have Atoh1occupany (**C**) Plot demonstrating the number and proportion of Atoh1 bound ATAC-seq sites that interact their nearest promoter (263/1854), their nearest promoter and others (542/1854) and other promoters (1049/1854). (**D**) Genome tracks (chr9:90,467,566-92,242,745) showing the PIFs proximal to the *Zic1* promoter. Promoter proximal PIFs are designated by arcs (grey for any PIF with significant CHiCAGO score ≥5, pink for PIFs that also contain an ATAC peak that overlaps with CHD7 binding signal, orange for PIFs that contain an ATAC peak that overlaps with Atoh1 ChIP peak and blue for a PIF that contains an ATAC peak overlapping with both a CHD7 and Atoh1 binding signals). The interval track below uses the same colour coding as the arcs (with the addition of black regions to show the captured promoter fragment) to show the width and location of the PIFs. A representative male ATAC signal track for WT (green) GCps are shown above the ChIP-seq signal tracks for Atoh1 (orange), CHD7 (grey), H3K27ac (pink) and H3K4me1 (dark blue).

Klisch et al. previously proposed a set of 601 Atoh1-regulated genes (P < 0.01) from an integrative analysis using linear DNA distances to link Atoh1, H3K4me1 and H3K4me3 with differential RNA expression in *Atoh1*-deficient GCps. We confirmed the interaction of 102 of these genes in our set of 577 Atoh1-bound genes derived from the pcHi-C analysis. These included genes with established roles in GCp development such as *Pax6*, *Neurod1*, *Neurod6*, *Nhlh1/2*, *Ccnd1*, *Ccnd2*, *Cxcr4* and *Sema7a*. By using pcHi-C we identified 475 additional Atoh1-regulated target genes, including *Zic1* and *Zic4*, and assigned these to their cognate enhancers (**Fig. 4D**; See **Suppl. Table 10** for the complete data).

### Atoh1 is recruited to the majority of CHD7-regulated enhancers

Having identified CHD7- and Atoh1-regulated enhancers, we next asked to what extent the CHD7- and Atoh1-regulated enhancers overlapped. We found that 829/1132 (73%) of CHD7-bound accessible regions also had Atoh1 recruited in GCps (**Fig.5A**). CHD7-bound sites without Atoh1 exhibited no change in chromatin accessibility in *Chd7*-deficient GCps in the majority of cases (290/303 = 96%), while 197/829 (24%) CHD7-Atoh1 co-occupied regions showed altered chromatin accessibility upon *Chd7* deletion. Thus, the majority of regions directly regulated by CHD7 are co-occupied by Atoh1 and chromatin accessibility changes upon CHD7 depletion appear to be strongly biased against regions without Atoh1. 197 Atoh1-CHD7 co-regulated enhancers could be assigned to 22 genes that were differentially expressed in both Atoh1- and *Chd7*-deficient GCps (overlap greater than expected by chance, *P=6.9x10-7,* One-way Fisher’s exact-test) (**Suppl. Table 11**). Several of these 22 genes regulated by CHD7 and Atoh1 have established critical roles in cerebellar development, including *Neurod2*, *Pax6* and *Gli2* (Corrales *et al*, 2004; Pieper *et al*, 2019; Swanson & Goldowitz, 2011) (**Fig. 5B**). As an example, our analysis identifies several potential Atoh1-CHD7 co-regulated *Gli2* enhancers (**Fig. 5C**) Pathway enrichment analysis of the 22 genes compared to all genes that were expressed in GCps shows a significant enrichment of terms: Hypoplasia of the pons (HP:0012110 *P*=0.006) and Abnormal pons morphology (HP:0007361 *P*=0.016) from human phenotype ontology, due to the presence of *Reln*, *Dcc*, (Rad et al, 2019) and *Gli2* (**Suppl Table 12**) (Corrales *et al*, 2006; Hong *et al*, 2000; Marsh *et al*, 2018).

**Figure 5.**
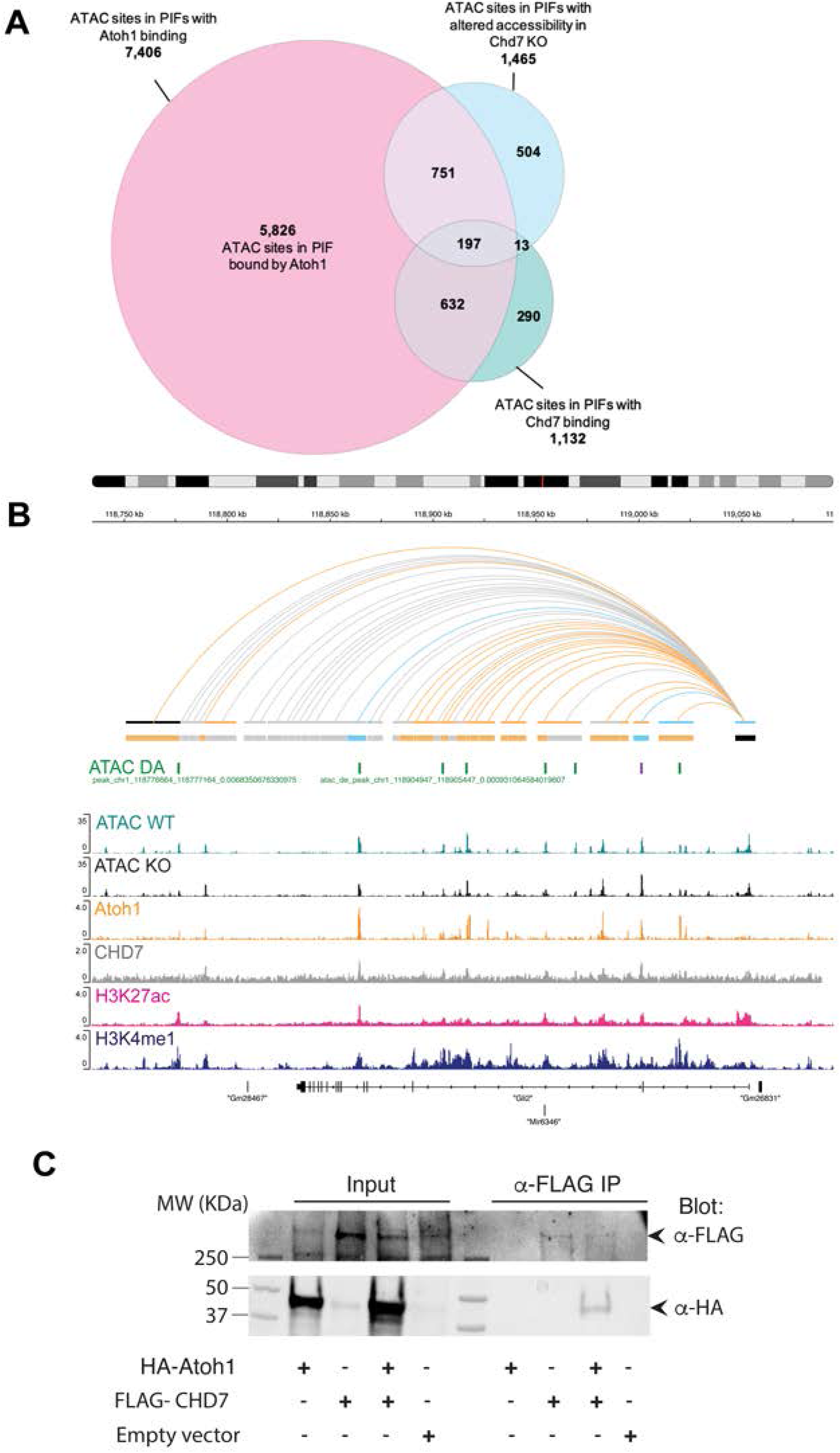
Atoh1 is recruited to the majority of CHD7-regulated enhancers. **(A)**Venn diagram demonstrating the total number of accessible chromatin sites in PIFs that also have Atoh1 recruitment (pink).The number of genomic locations that demonstrate altered accessibility in *Chd7*-deficient GCps (light blue), 751/1465 of which were in PIFs that demonstrate Atoh1 recruitment. The number of ATAC sites with altered accessibility in *Chd7*-deficient GCps that also exhibit CHD7 recruitment in PIFs (light green), 632/1132 of which were in PIFs that also demonstrate Atoh1 recruitment. Finally, 197 PIFs were identified that exhibit differential accessibility in *Chd7*-deficient GCps and to which both Atoh1 and CHD7 are recruited. **(B)** Genome tracks at the *Gli2* locus (chr1:118,717,920-119,121,120) showing the location of three putative, promoter-proximal PIFs designated by arcs (grey for any PIF with significant CHiCAGO score ≥5, orange for PIFs that also contain an ATAC peak that overlaps with Atoh1 binding signal and blue for PIFs that contain accessible regions that overlap with regions of Atoh1 and Chd7 binding). Regions of differential chromatin accessibility (p-adj < 0.05) are displayed in the interval track (ATAC DA) showing 500bp regions of decreased (green) and increased (purple) accessibility in *Chd7*-deficient GCps between the WT ATAC (n=3) and *Chd7*-deficient (KO) ATAC signal (n=3). A representative ATAC signal track for WT (green) and CHD7 KO (black) GCps are shown above the ChIPseq signal tracks for Atoh1(orange), CHD7 (grey), H3K27ac (pink) and H3K4me1 (dark blue). **(C)** Co-immunoprecipitation of ATOH1 and CHD7 in HEK293T cells, transiently transfected with FLAG-tagged CHD7, HA-tagged Atoh1, both or empty vector (EV). CHD7 was immunoprecipitated with anti-FLAG followed by western blot against FLAG and HA. Note the co-immunoprecipitation of HA-tagged ATOH1 upon FLAG-CHD7 immunoprecipitation.

These findings suggested that Atoh1 and CHD7 are frequently recruited to the same enhancers in GCps. We therefore tested if Atoh1 and CHD7 complexes might interact directly. Indeed, co-immunoprecipitation experiments revealed that CHD7 and Atoh1 interact with each other (**Fig. 5C**).

Together, these findings suggest that CHD7 cooperates closely with Atoh1 in GCps, akin to other lineage-specific transcription factors. However, we also note that the majority of putative regulatory elements with Atoh1 recruitment in GCps did not appear to be regulated by CHD7 (**Fig. 4A**), suggesting that Atoh1 may regulate other enhancers and genes in concert with other transcription factors and chromatin remodelling factors.

## Discussion

Understanding the mechanisms whereby distal enhancers regulate developmental processes remains a significant challenge, partly due to the fact that these mechanisms operate over long genomic distances. Here we used pcHi-C to identify distal regulatory elements of genes expressed in cerebellar GCps during early postnatal development,enabling the mechanistic interrogation of *bona fide* enhancer-promoter interactions in GCps. By integrating pcHi-C, histone ChIP-seq and ATAC-seq data, we identified 46,428 promoter-enhancer interactions in cerebellar GCps, including 22,797 potential distal regulatory regions for 12,905 genes. We illustrate the utility of this data by identifying the genes regulated by hindbrain VISTA enhancers in GCps, and identifying genes regulated by CHD7- and Atoh1-regulated enhancers. Our analysis identified CHD7-Atoh1 co-localisation at many enhancer regions, and together with a demonstration that Atoh1 and CHD7 can interact in co-immunoprecipitation experiments, suggest that Atoh1 and CHD7 function together to control gene expression in GCps.

We previously reported a significant downregulation of the *Reln* gene in *Chd7*-deficient GCps and showed that *Reln* downregulation was partly responsible for the cerebellar hypoplasia in these mutants. We identified putative enhancer regions 127kb upstream and in exon 8 of *Reln*, but without further evidence, it was not known if these regions were *Reln* enhancers (Whittaker *et al*., 2017b). Using our pcHi-C data, we found that *Reln* is indeed a direct target of CHD7, but that CHD7-regulated *Reln* enhancers are located in *Reln* intronic regions 249kb, 395kb and 396kb downstream of the *Reln* promoter (see **Fig. 3C**).

In addition to confirming that *Reln* is indeed a direct target of CHD7, we identified 62 other direct CHD7 target genes with their cognate distal enhancers. These direct targets included several genes with known functions in GCps. Perhaps surprisingly, two of these genes, *Gli2* and *Ccnd1*, with known functions as mediators of SHH-induced GCp proliferation (Corrales, Rocco et al. 2004, Pogoriler, Millen et al. 2006), were upregulated in *Chd7*-deficient GCps. Upregulation of these genes are expected to lead to cerebellar hyperplasia, rather than the observed hypoplasia. One possible explanation is that the expression of other genes might counter-act the effects of *Gli2* and *Ccnd1* upregulation by having more pronounced effects on GCp proliferation. Another intriguing possibility is that the increased expression of *Gli2*, which is also known to mediate the SHH-induced foliation in the developing cerebellum (Corrales, Rocco et al. 2004, Corrales, Blaess et al. 2006), might mediate some of the cellular changes that underlie the polymicrogyria phenotype reported in these mutants (Reddy, Majidi et al. 2021).

We compared our pcHi-C data with data from Reddy et al., who performed Hi-C from the anterior cerebellum of P4 mice. This study identified 5,174 enhancer-promoter (E-P) interactions, compared to the 46,428 promoter-accessible PIF regions identified in the present study, demonstrating the advantage of promoter capture in enriching for E-P interactions. Reddy et al. identified long-range interacting regions for 1207 genes. A substantial portion of these genes (843/1207=82%) are also represented in our pcHi-C dataset (**Suppl. Fig. 3A, Suppl Table 13**). Comparing the E-P interactions of these 843 genes between these two studies found the same E-P interactions for 38% (**Suppl. Fig. 3B**) This relatively low overlap can be explained by two differences: 53% of the 3229 interactions unique to the Reddy data are not detectable in the pCHi-C data due to the removal of bait-to-bait interactions during our data analysis or the absence of probes to capture gene promoters (**Suppl. Fig. 3B**). For the remaining interactions identified by Reddy et al, we identify other distal interacting regions, possibly due to E-P interactions in cell types other than GCps being detected in the Reddy study. In addition to experimental differences, there are also notable differences in analysis strategies. Reddy et al. binned the Hi-C data into 10kb regions to identify interacting regions and subsequently used chromatin marks to identify possible enhancer and promoter regions within these large regions. By contrast, we have used the pCHi-C and CHiCAGO algorithm to identify individual HindIII restriction fragments and prioritised those that have accessible regions within them for integration with chromatin modifications.

We demonstrated the utility of pcHi-C to assign genes to distal regulatory elements. Most importantly, we found that 80% of the putative enhancers we identify bypass their neighbouring genes to interact with a more distal gene. This is consistent with a recent study by Chen et al. who found 61% of developmental enhancers bypassing their closest genes (Chen *et al*., 2024). Thus, inferring enhancer-gene regulatory relationships based on linear proximity on the genome in cis is unreliable. Similarly, we found that 77% of VISTA hindbrain enhancers were incompletely annotated. However, it should be noted that unctional studies are still required to confirm enhancer-gene regulatory relationships inferred from pcHi-C data.

Identifying CHD7-regulated enhancers is challenging, mostly due to the low quality of available CHD7 ChIP-seq data. However, integrating CHD7 ChIP-seq data with ATAC-seq accessibility, histone modification ChIP-seq and pcHi-C data has allowed us to identify a subset of enhancers that are most likely directly regulated by CHD7. However, given these technical limitations, we would be hesitant to conclude from the present data that the majority of chromatin accessibility changes in enhancers in *Chd7*-deficient GCps are indirect, as suggested by the data in Fig. 3A.

Previous studies have shown that CHD7 cooperates with lineage-specific transcription factors. This suggests that CHD7 is recruited to specific sequence motifs chromatin via lineage-specific transcription factors. In addition to already reported interactions with Sox2 in neural and oligodendrocyte progenitors (Engelen, Akinci et al. 2011, Doi, Ogata et al. 2017) , Sox10 in oligodendrocyte progenitors (He, Marie et al. 2016), Runx1 in haematopietic stem and progenitor cells (Zhen, Kwon et al. 2017, Hsu, Huang et al. 2020), and Isl1 in secondary heart field (Stathopoulou, Wang et al. 2023), we show here that CHD7 interacts with Atoh1 and is recruited with Atoh1 to enhancers in cerebellar GCps. Based on studies in other cellular contexts, it seems likely that Atoh1 recruits CHD7 to enhancers in GCps, where CHD7 remodels chromatin, although cooperative recruitment of CHD7 and Atoh1 and other non-remodelling functions of CHD7 cannot be excluded without further experimentation. It also seems likely that CHD7 might control later stages of GCp differentiation by interacting with other neurogenic bHLH transcription factors like Neurod1.

Our analysis confirmed that Atoh1 is preferentially recruited to distal enhancers, marked by H3K4me1 (Klisch *et al*., 2011). In our analysis, 10,0505 ATAC-seq peaks with Atoh1 were distal intergenic, 7,837 intronic, with only 3,602 located <3kb of the promoter (<600 each further away from the promoter, exonic or in 5’ or 3’ UTRs). We were able to identify 679 gene promoters that were in close proximity to Atoh1-bound distal regulatory elements, 102 of which were proposed by Klisch et al. and 475 new Atoh1-regulated target genes.

Although the present study represents a significant advancement in our ability to assign distal regulatory elements to their cognate gene promoters in primary GCps, our study and analysis have certain limitations. First, although pcHi-C significantly enriches for promoter-enhancer interactions, and we sequenced our libraries at considerable depth, the fact that our replicate libraries showed incomplete (57-65%) overlap, suggests that coverage is still limited. Additional replicates would enable further confidence in results and potentially increase coverage of under sampled regions. A similar limitation has to be recognised for other datasets e.g. ChIP-seq and ATAC-seq. For instance, our data suggests that the majority of Atoh1-bound enhancers are not regulated by CHD7 (**Fig. 5A**). However, it remains possible that more of these Atoh1-bound elements are regulated by CHD7 but our ability to detect these by CHD7 ChIP-seq and differential ATAC-seq analyses are limited. However, whilst individual datasets each have their own have limitations, by combining and intersecting datasets from multiple sources we have built a robust set of distal regulatory elements supported by multiple datasets.

## Materials and methods

### Animals

Mice were bred and maintained according to Home Office regulations in New Hunt’s House Biological Service Unit, King’s College London. C57BL/6J animals were supplied from internal breeding stocks at the New Hunt’s House Biological Service Unit, King’s College London.

### Cerebellar granule cell progenitor purification

GCp purification was performed as previously described (Hatten, 1985; Whittaker *et al*., 2017b). Cerebella were dissected at P7 in DPBS (Gibco 14190144, 0.0001% phenol red) ensuring to remove the meninges. Cerebella were cut into small pieces and dissociated using Papain I (100U; Worthington) in DPBS with 200μl DNase I (Sigma, D4527) and 2mg L-Cysteine (Sigma, C1276), for 30 minutes at 37°C. Subsequently, GCps were isolated by a two-step Percoll (Sigma, P4937) gradient (35%/65%). The GCps on the 35%/65% interface were taken for further analyses.

### Luciferase assays

Two PIFs identified by pCHi-C were PCR amplified from mouse genomic DNA using the following primer sequences that contained KpnI and XhoI restriction enzyme sites for cloning and 3 additional nucleotides prior to the restriction enzyme recognition site to enhance cutting efficiency:

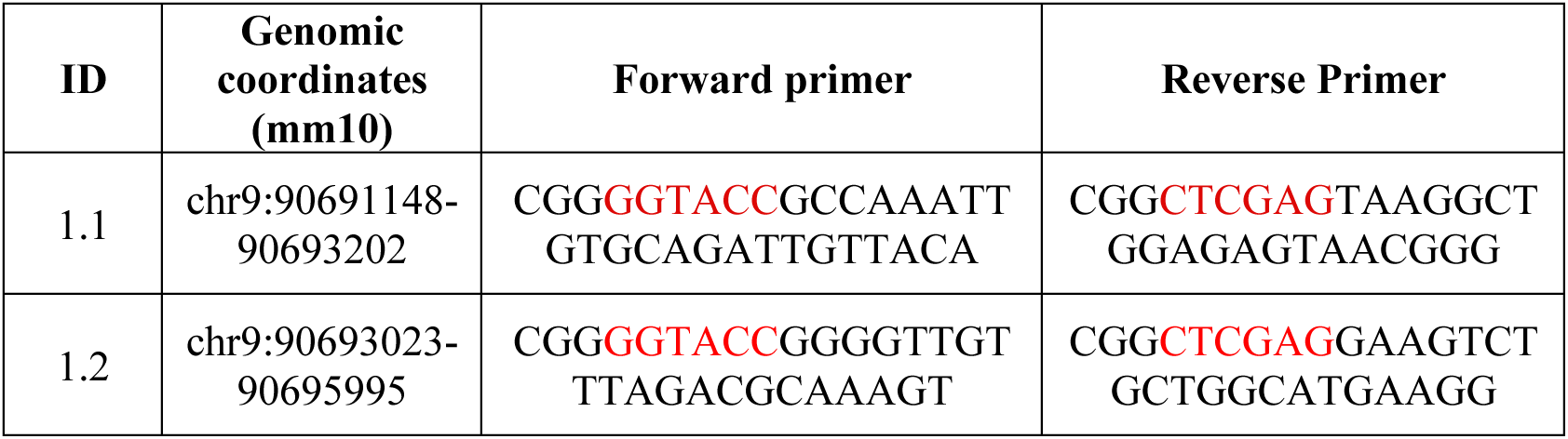

PCR products were run on a 0.8% agarose gel and extracted using the Monarch DNA Gel Extraction Kit, per manufacturer’s instructions. DNA was digested with KpnI (NEB), at 37°C for 15 minutes. The reaction was cleaned up using the Monarch PCR and DNA Cleanup Kit and DNA eluted in 6µl elution buffer. Subsequently, the DNA was digested with XhoI (NEB) at 37°C for 15 minutes. XhoI was heat-inactivated at 65°C for 20 minutes and the reaction was cleaned-up using the Monarch PCR and DNA Cleanup Kit. DNA was eluted in 6µl. 1µg of pGL3 was digested with KpnI and XhoI as described above and dephosphorylated with rSAP at 37°C for 30 minutes, followed by a 5 minute heat-inactivation step at 65°C. Subsequently, the vector was run on a 0.8% gel, extracted using the Monarch DNA Gel Extraction Kit and eluted in 10µl elution buffer. A ligation reaction using the NEB T4 DNA ligase, as per manufacturer’s instructions. A vector only ligation was always run alongside each ligation, as an internal control for vector self-ligation. 100ng of vector was used in ligations and typically a 3:1 molar excess of the insert was added into the reaction, as calculated using the NEBioCalculator Tool. Ligations were performed at 16°C in the Eppendorf Mastercycler Gradient o/n. Subsequently, the T4 DNA ligase was heat-inactivated at 65°C for 15 minutes and the ligation reactions stored at -20°C until further use. Ligation reactions were transformed into NEB 5-alpha Competent E.coli (High Efficiency), as per manufacturer’s instructions. 150µl of the reactions were spread on pre-warmed ampicillin (100µg/ml final concentration) LB Agar plates and incubated at 37°C o/n. Ligations were deemed successful when the vector only ligation ampicillin plate contained <10% colonies compared to experimental plates. Single colonies from experimental plates were picked and grown in mini-cultures of 6-8ml LB (100µg/ml ampicillin) o/n. To screen colonies for successful integration of the insert, 1ml of the mini-cultures were used and plasmid DNA isolated using the Monarch Plasmid Miniprep Kit, per manufacturer’s instruction. mini-prep DNA was used for a diagnostic digest, using KpnI and XhoI in the same 25µl reaction, and DNA run on a 0.8% gel. Clones that contained bands of the correct size for both the vector and insert were sent for Sanger sequencing. Plasmid DNA of reactions demonstrating correct integration of the insert were extracted using the kit-free mini-prep protocol from Addgene.

To introduce plasmid DNA into SHH-NPDs, the SG Cell Line 4D-Nucleofector X Kit was used. 1x10^5^ cells were resuspended in 20µl of SG nucleofection solution and 400ng of each of the desired plasmid(s) DNA was added. DNA volume never exceeded >10% of the final sample volume. This mix was rapidly added to 1 well in the 16-well cuvette and the cells were nucleofected using the CM138 setting on the X-unit of the 4D Nucleofector unit. Following nucleofection, the cells were left to recover for 10 minutes at RT. Subsequently, 80µl of pre-warmed whole cell media was added to the cuvette well harbouring the nucleofected cells and this mix was plated in 100µl of pre-warmed media in a flat bottom Greiner 96 well plate.

Luciferase assays were performed 36-48h later using the Dual-Glo Luciferase Assay System in a flat bottom Greiner 96 well plate on the GloMax 96 Microplate luminometer. The Renilla vector was always used in each well as an internal transfection control and the empty pGL3 vector as the experimental control. Experimental (firefly, pGL3) luminescence readings were divided by Renilla luminescence readings for each well tested, to gain relative luminescence values. The relative luminescence values for experimental conditions were divided by the average of the relative luminescence for the experimental control (empty pGL3) to calculate normalized luminescence.

### Co-immunoprecipitation

Human embryonic kidney 293T (HEK293T) cells were cultured in Dulbecco’s modified Eagle’s medium (DMEM; Sigma) with 10% fetal bovine serum (FBS; Thermo Fisher Scientific), 1% l-glutamine (Gibco) and 1% penicillin- streptomycin (Gibco). Cells were maintained in a 5% CO_2_ humidified incubator at 37°C. Twenty- four hours prior to transfection, cells were plated into a 10cm dish in media without penicillin-streptomycin. After 24 hours, when 60-70% confluent, cells were transiently transfected with FLAG- tagged CHD7 inserted into a pcDNA3.1(+) vector (Bajpai *et al*, 2010), HA- tagged Atoh1 inserted in pHM6 (Krizhanovsky *et al*, 2006) or control (empty vector) expression constructs in OptiMEM (Thermo Fisher Scientific) using Lipofectamine (Invitrogen), according to manufacturer’s instructions. Cells were cultured for 60 hours following transfection. Cells were collected in ice cold PBS, pH 7.4 (Thermo Fisher Scientific) with protease inhibitors (cOmplete, Sigma-Aldrich) using a cell scraper and pelleted by centrifugation (1100rpm, 4°C for 10 minutes). Nuclear extraction was performed as previously described (Dignam *et al*, 1983). In brief, pelleted cells were resuspended in buffer A (10mM Hepes-KOH pH 7.6, 1.5mM MgCl2, 10mM KCl) with protease inhibitors and were incubated on ice for 10 minutes to allow cell lysis. The nuclei were lysed by incubation in buffer C (20mM Hepes-KOH pH 7.6, 20% glycerol, 420mM NaCl, 1.5mM MgCl2, 0.2mM EDTA) with protease inhibitors at 4°C for 30 minutes. The nuclei and protein precipitates were removed by centrifugation and the supernatant containing the nuclear extract was transferred to a new eppendorf tube. After protein quantification with a Bradford assay, samples were diluted with C-0 buffer (20mM Hepes-KOH pH 7.6, 20% glycerol, 1.5mM MgCl2, 0.2mM EDTA) to a final salt concentration of 200mM and 300µg of protein from each sample used for immunoprecipitation. Magnetic flag beads (anti- FLAG M2 magnetic beads, Millipore) were added to each sample for immunoprecipitation and incubated for 4 hours at 4°C rotating. Samples were washed with C-100 buffer (20mM Hepes- KOH pH7.6, 20% glycerol, 200mM NaCl, 1.5mM MgCl2, 0.2mM EDTA, 0.02% NP-40 (Sigma) and eluted in SDS loading buffer with beta mercaptoethanol. Co-immunoprecipitation was evaluated by western blot using anti- FLAG (F3165, Sigma-Aldrich, 1:5000) and anti- HA antibodies (C29F4, Cell signalling, 1:1000).

### pcHi-C

PCHi-C was carried out essentially as described (Villiers *et al*, 2023). 3-4 x10^7^ isolated GCps were fixed in a 2% final concentration of formaldehyde for 10 min at room temperature. The reaction was quenched, the cells resuspended in ice-cold lysis buffer and incubated for 30 min at 4°C. Approximately 1 x 10^7^ GCps were incubated overnight with 1500U of HindIII (NEB, R0104T) at 37°C. DNA ends were labelled with biotin-14–dATP (Life Technologies, 19524-016) in a Klenow (NEB, M0210L) end-filling reaction. Blunt-end ligation using T4 DNA ligase (Life Technologies, 15224-025) was performed at 16°C overnight, followed by an overnight proteinase K digestion (20mg/ml;Roche, 3115879001) at 65°C to remove protein. DNA was purified by a double phenol-chloroform extraction and quantified using the Qubit broad range assay (Qubit). Subsequently, biotin was removed from non-ligated DNA ends in a T4 polymerase (NEB, M0203L) reaction and again subjected to phenol-chloroform purification. To obtain an average of 400bp fragments, DNA was sheared using the Covaris Focused ultra-sonicator, according to the manufacturers’ instructions. The sheared DNA was end-repaired, adenine tailed and double size selected employing AMPure XP beads (Beckman Coulter Ampure XP beads, A63881) to isolate DNA ranging from 250 to 550 bp in size. Ligation junctions were isolated using Dynabeads MyOne Steptavidin T1 beads (Life Technologies, 65601). DNA was ligated to paired-end adaptors (Illumina) and amplified using PE PCR 1.0 and PE PCR 2.0 primers (Illumina) for 6 cycles. For promoter capture, the Hi-C library was hybridised to 39,684 custom-designed biotinylated RNA probes “baits’’, using SureSelect Target Enrichment kit (Agilent Technologies) according to the manufacturer’s instructions. The baits were originally designed to cover 22,047 promoters of mm9 genome (**Suppl. table 1)** as described (Yun *et al*, 2021b). Junctions that had hybridised to RNA baits were pulled down using Dynabeads MyOne Streptavidin T1 (Life Technologies, 65601). The library was amplified using PE PCR 1.0 and PE PCR 2.0 primers (Illumina). Two replicate libraries were prepared as described and sent for paired-end sequencing (75bp) on a HiSeq 4000 platform to a depth of 675,941,525 and 589,050,886 read pairs for the two replicates. Each replicate required two sequencing runs to reach sufficient sequencing depth.

### pcHi-C data processing and interaction calling

For each pcHiC replicate Fastq files from individual sequencing runs were combined and each replicate was processed separately using the HiCUP pipeline version 0.5.8 (Wingett *et al*., 2015). Briefly, a HindIII restriction fragment map for the mm10 genome (chromosomes 1-19,X,Y & M only, random and ChrUN contigs were removed) was obtained by running the hicup_digester tool with the options ‘ -z --re1 A^AGCTT,HindIII’. Read mapping was performed using Bowtie 1.1.2 (Langmead *et al*, 2009) to the mm10 genome (chromosomes 1-19,X,Y & M only) and default parameters were used to remove reads resulting from ligation artefacts and PCR duplication to obtain valid di-tags to take forward to CHiCAGO analysis.

Prior to interaction calling mm9 probe coordinates were converted to mm10 using UCSCliftover (Hinrichs *et al*, 2006) and HindIII digested mm10 genome fragments were deemed to be pulled down by the probes and assigned as ‘bait’ regions if they overlapped the probe fragments by >= 110bp. This resulted in a Baitmap input file for CHiCAGO analysis (Cairns *et al*., 2016) of 39,671 probes targeting 22,449 unique HindIII restriction fragments. All HindIII restriction fragments were annotated by intersecting fragments with gene and transcript annotations from Ensembl version 81 using pybedtools (0.7.10). Bait names were reassigned based on their overlap with transcript TSS, or if no TSS was present in fragment the name was assigned as overlapping the gene body of any transcripts the bait overlapped, and if no overlap with any transcripts was detected the bait name was updated to indicate this along as well as a reference to the regions it was intended to target. CHiCAGO design files and chinput files for each replicate were generated using default options with the chicagoTools scripts from the CHiCAGO package version 1.12.0. Both replicates were provided to the RunChicago.R script (R version 3.6.1) to produce a ‘superset’. Bait-PIF interactions were identified using default parameters and deemed significant with a CHiCAGO interaction score >=5. RunChicago.R was run 10 times, and the output with the fewest number of significant bait-PIF interactions was taken forward and filtered to remove trans chromsomal PIFs, bait-to-bait PIFs and interactions where the baits did not overlap a TSS of a transcript from the Ensembl 81 gtf with a ‘protein_coding’ gene_biotype. Summary metrics (cumulative distance percentiles) were calculated using Python & plotted using R.

It should be noted the number of reads needed to score an interaction by the CHiCAGO algorithm depends on the both the distance between a PIF and the promoter, modelled using a distance-dependent component that accounts for decay of contact frequence with genomic distance, and a component that models how the sequence or other technical artifacts might influence the capture bias of some sequences compared to others. For each promoter a background model is generated of the expected number of reads that would be captured based on the above considerations and if the number of reads for that region exceeds this background model by a certain threshold the interaction is deemed significant using a p-value like score. In practice this means that regions further from the promoter will often require fewer reads to signify a significant interaction compared to regions that are much closer to the promoter. The significant PIFs in the dataset are all evidenced by a minimum of 3 reads in at least one biological replicate.

### ChIP-Seq for H3K4me1, H3K4me3 and Atoh1

ChIP-seq data (Klisch *et al*., 2011) were downloaded from ENA (PRJNA128899) trimmed for Truseq2 adapter sequences using Trimmomatic (v0.36) in single end ILLUMINACLIP mode with default parameters apart from setting mismatches to 1 and MINLENGTH:24. Paired-end H3K27ac & CHD7 ChIP-seq data from P7 GCps (Feng *et al*., 2017) were downloaded from ENA (PRJNA309175) and adapters were removed using Trimmomatic PE ILLUMINACLIP with the settings 1:30:10:1:True LEADING:3 TRAILING:3 SLIDINGWINDOW:4:15 MINLEN:36. Reads were mapped to the mm10 genomes using bwa aln v0.7.17 (Bolger *et al*, 2014). Unmapped and secondary alignment reads were filtered using SAMTools (Danecek *et al*, 2021) and BEDTools intersect (Quinlan & Hall, 2010) was used to remove those in encode blacklisted regions (Amemiya *et al*, 2019). Duplicate reads were removed using PICARD MarkDuplicates (v2.18.0, http://broadinstitute.github.io/picard). Replicates were then combined where available and peakcalling performed using MACS2 (Zhang *et al*, 2008) with parameters callpeak --bdg --SPMR --gsize mm --call summits –tsize against a GCps ChIP-seq input control where available. H3K4me1 peakcalling used the additional --broad parameter. Bigwig ’treatment’ coverage tracks produced by MACS2 were used for visualisation.

### ATAC-seq

ATAC-seq libraries were the same as those prepared for our previously published manuscript (Whittaker, Riegman et al. 2017) (GSE90466) and reanalysed for this study. Reads were trimmed to remove adapters using Trimmomatic v0.36.6 with the parameters ‘-phred33 NexteraPE-PE.fa:1:30:10:1:True LEADING:3 TRAILING:3 SLIDINGWINDOW:4:15 MINLEN:30’, and aligned to mm10 (chr1-19,M,X & Y) with Bowtie2 v2.3.4.1 (Langmead & Salzberg, 2012) with default parameters plus -X2000. Aligned reads were filtered using PICARD v2.18.0 MarkDuplicates ‘ASSUME_SORTED=true REMOVE_DUPLICATES=true VALIDATION_STRINGENCY=SILENT’ to remove duplicates and samtools v1.9 view to remove reads that were unmapped (-F 4), secondary alignments (-F 0x100), unpaired (-f1 -f 2), low quality with a MAPQ < 30 (-q 30) and those on chrM.

A mm10 combined blacklist of regions with high similarity to mitochondrial reads and high signal artifacts identified by ENCODE was downloaded from https://github.com/buenrostrolab/mitoblacklist/tree/master/combinedBlacklist (Buenrostro *et al*, 2015), reads overlapping these regions were removed using bedtools v2.27.1 pairToBed ‘-f 0.00000010 -type neither’.

The master peaklist was generated from WT samples by downsampling each sample to 43,355,200 reads, then combining all into to a single bam file and calling peaks using MACS2 callpeak (v2.1.1) and the following arguments ‘--format=BAMPE --bdg --SPMR --call-summits --keep-dup all --gsize mm --tsize 58’. Summit regions were extended +/- 250bp to create 500bp wide regions. When regions overlapped those with the highest -log10 p-value were retained, producing a final peak list of 112,341 non-overlapping 500bp regions. For differential accessibility analysis reads in WT peaks list were counted for the 3x WT and 3x *Chd7*-deficient samples (not downsampled) using deepTools multiBamSummary --outRawCounts 3.1.3 (Ramirez *et al*, 2016). Differential accessibility analysis was performed using DESeq2 (v 1.22.1) (Love *et al*, 2014). Whilst all peaks were tested for differential accessibility, size factors were calculated using only promoter peaks of genes whose RNA was not differentially expressed in the *Chd7*-deficient GCps. Homer annotatePeaks.pl (v4.10) (Heinz *et al*, 2010) was used to annotate peak location of promoters based on Ensembl 81 GTF file and those assigned as ’promoter-TSS’ were selected for size factor calculation if their nearest gene ID had a p-adjusted value > 0.95 in the RNA-seq differential expression (see RNA-seq section for details). The DESeq2 design used a model controlling for sample sex (design=∼sex + genotype) and multiple testing correction was applied using the Independent Hypothesis Weighting package (version 1.10.1) (Ignatiadis *et al*, 2016) where alpha = 0.05. Log-fold change values were calculated using the ’normal’ shrinkage method. Peaks with a p-adjusted value < 0.05 were deemed to be differentially accessible in the *Chd7*-deficient GCps.

### ATACSeq Coverage Tracks

ATAC-seq bigwigs for individual libraries were generated using deepTools bamCoverage (v3.1.3), using the parameters ‘--binSize 1 --extendReads --scaleFactor’ the scaling factor for each sample was computed using 1/(total reads in peaks on chromosomes 1-19)*1^6^. WiggleTools (v1.2.3) (Zerbino *et al*, 2014) write_bg was then used to create coverage tracks representing the median coverage of the replicates and coverted to bigwig files using ucsc bedGraphToBigWig (v377), (Kent *et al*, 2010).

### RNA-seq

RNA-seq libraries were the same as those prepared for our previously published manuscript (Whittaker *et al*., 2017b) (GSE90466) and reanalysed for this study. Reads were quantified using Kallisto version 0.43.0 (Bray *et al*, 2016). A Kallisto index was generated using -k 31 and a fasta file generated for exonic regions of transcripts from Ensembl 81 annotations where gene_biotype = “protein coding”. Quantification was performed using ‘Kallisto quant --bias -t 1 --bootstrap-samples=100’. TPM values were transformed to gene level counts using tximport (1.6.0) (Soneson *et al*, 2015) and differential expression was performed at the gene level using DESeq2 (1.18.1) with the model design (∼replicate + group). Multiple testing correction was applied using Independent Hypothesis Weighting (IHW) with a default alpha of 0.1, and genes deemed differentially expressed if p-adjusted value of < 0.05. Log-fold change values were generated using ’normal’ Bayesian shrinkage method. The RNA-seq data from Klisch et al. (2011) was analysed as above, but due to an imbalance of sex-specific genes between the WT and KO samples chr X&Y were excluded from the differential expression analysis. Expression heatmaps were generated using pheatmap (https://CRAN.R-project.org/package=pheatmap) from log2(normalised counts +1) and scaled by row. Genes were ordered via log-fold change.

### RNA-seq Coverage Tracks

Coverage tracks were created by trimming reads to remove adapters using Trimmomatic (0.32) with the parameters ‘-phred33 ILLUMINACLIP:TruSeq2-PE.fa:1:30:10:8:true LEADING:3 TRAILING:3 SLIDINGWINDOW:4:15 MINLEN:36’. Alignment was performed using STAR 2.4.2a ‘--runMode alignReads --outSAMstrandField intronMotif --outSAMunmapped Within --outFilterType BySJout --outFilterMismatchNmax 999 --outFilterMismatchNoverLmax 0.04 --outFilterIntronMotifs RemoveNoncanonicalUnannotated’ (Dobin *et al*, 2013). Individual bigwig files were normalised using sizeFactors generated from DESeq2 analysis using deeptools 3.4.3 bamCoverage --binSize1, --scaleFactor 1/sizeFactor and replicates were then combined using wiggletools write_bg (1.2.3) and bedGraphToBigWig (version 377) to generate mean coverage tracks.

### Publicly available datasets used

H3K27ac and CHD7 ChIP-seq data from P7 GCps from (Feng *et al*., 2017) (GSE93741), ATOH1 ChIP-seq at P5 GCps and E18.5 rhombic lip RNA-seq with or without inducible *Atoh1* gene deletion in the Atoh1+ lineage (Klisch *et al*., 2011) (GSE22111) were downloaded as raw sequencing files from the European Nucleotide Archive and processed as described above.

Locations of all 140 mouse loci- Mouse hindbrain enhancers were downloaded from the Vista Enhancer Browser (Visel *et al*., 2007) and converted to mm10 coordinates using UCSC liftOver (Hinrichs *et al*., 2006) for intersection with PIFs.

Ensembl heterochromatin features were obtained from the Ensembl (version 81) Regulatory build segmentation file for brain E14.5 and filtered for regions predicted to be heterochromatin (Zerbino *et al*, 2016).

Postnatal day 6-9 FANTOM5 Cerebellum Enhancers (N06 & N09) were curated by selecting the libraries for postnatal day 6 and postnatal day 9 mouse cerebellum samples (n=3 for each timepoint) from F5.mm10.enhancers.expression.usage.matrix.gz (https://doi.org/10.5281/zenodo.1411211). 818 regions that were present in at least 2 of the libraries were retained for CHiCAGO enrichment analysis.

### Integration and Analysis

#### PCHiC analysis and enrichment of features

Absolute distances between interactions were calculated from the mid-point of each restriction fragment. Enrichment of PIFs for H3K27ac, H4K4me1, FANTOM5 enhancers and encode heterochromatic regions analysis was performed using CHiCAGO using all interacting fragments with protein-coding baits, overlap of randomly sampled distance-matched subsets was performed 100 times and using default settings. Weighted mean of scaled read counts were calculated using CHiCAGO.

#### Nearest gene Analysis

Locations of PIFs fragments significantly interacting with protein coding bait regions were annotated using the ChIPseeker package (v 1.22) (Yu *et al*, 2015) and assigned to their nearest protein coding gene using a custom transcript database generated by txdb bioconductor package after extracting protein coding genes (gene_biotype: ‘protein_coding’) from the mm10 Ensembl (version 81) annotation gtf file. These nearest gene annotations were compared with the annotations of promoters linked to all PIFs, those containing VISTA enhancer fragments and those with Atoh1 bound ATAC-seq peak regions using custom python scripts.

#### Assessment of ATAC-seq peaks in PIFs

PIFs overlapping an ATAC-seq site by at least 1 bp (as determined using pybedtools intersect) were assigned as PIFs containing an ATAC-seq site. Distances from PIF without an ATAC-seq peak to a PIF with an ATAC-seq peak were calculated using pybedtools closest using default parameters. The PIF with the highest CHiCAGO score for each promoter bait fragment was selected and assigned the ‘lead PIF’ for that bait. The 10 HindIII fragments flanking either side of the ‘lead PIF’ were designated as being a PIF if an interaction of that fragment was found with any bait. The fragment was assigned as containing an ATAC-seq peak if it intersected any peak from the wild-type (WT) ATAC-seq peak list.

#### pcHi-C, RNASeq, ATAC-seq and ChIP-seq Integration

Probed HindIII fragments containing a TSS for a protein coding transcript were assigned to the expression levels and differential expression status in *Chd7* and *Atoh1* RNA-seq experiments of their corresponding protein_coding gene. Mean expression level in *Chd7 +/+* and *Atoh1* +/+ mice (TPM) were assigned to genes in baits and categorised into 5 categories: ’not expressed’, if TPM < 4 with the remaining expression values divided into quartiles and labelled as ’very_low’,’low’,’mid’,’high’ expression genes.

All WT ATAC-seq peaks were annotated in a binary fashion as to whether they intersected with CHD7 & Atoh1, H3K27ac, H3K4me1 and H3K4me3 ChIPSeq peaks using pybedtools intersect along with their differential accessibility status in *Chd7*-deficient mice. These annotated ATAC-seq sites were used to annotate all HindIII fragments to enable identification and assess properties of ATAC-seq sites contained within PIFs. ChIPSeeker (v 1.22) was used to annotate ATAC-seq locations as described above for PIFs. Combinations of Histone marks and locations were plotted using UPSetR and ggplot.

#### Motif Enrichment

Motif enrichment was performed using SEA from MEME suite (v5.5.5) (Bailey & Grant, 2021a) with the Mouse HOCOMCOCOv11 CORE database. 100bp fasta sequences were prepared with 50bp +/- from the summit of each WT ATAC-seq peaks. Enrichment of accessible peaks in PIFs was tested against a background of all ATAC-seq peaks, using default options. Where multiple motifs for the same DNA binding factor were recovered the most significant was taken through to further analysis. Significantly enriched motifs were filtered to remove motifs with (q-value <0.05) and motifs of genes with an expression level of TPM < 4 (i.e. ‘not expressed’) in the wildtype RNA-seq data. Motifs were annotated using TFClass (REF) in R and grouped by transcription factor ‘family’.

#### Gene and Pathway Enrichment

Over representation analysis of genes was performed using gprofiler2 v0.2.1 (Kolberg *et al*, 2023) using Ensembl ID’s and excluding electronically inferred annotations in R. For enrichment of genes with/without ATAC-seq sites in PIFs results were generated against a background gene-set of all protein coding genes. Results were filtered to only include GO:BP genesets containing between 200-400 terms (Biomart release 2024-01-17). Over representation analysis of 63 genes that were differentially expressed in *Chd7*-deficient GCps and had PIFs containing accessible regions that overlapped with CHD7 ChIP-seq peaks and were differentially accessible in ATAC-seq from *Chd7*-deficient GCps were compared against all protein-coding genes that were included in the differential expression analysis by DESeq2 analysis (i.e. with padj > 0), no post-analysis filtering was performed based on number of terms in gene set. Over representation analysis of genes differentially expressed in *Atoh1*-deficient cerebellar tissue and *Chd7*-deficient GCps was performed against a background of genes that were included in both differential expression analyses and gene-sets were not filtered post-analysis for number of terms in gene-sets.

#### Klisch significant targetome

601 genes identified in Atoh1 targetome (p<0.01) by Klisch et al. (2011) were obtained from Supplementary Table 5 of their paper. Ensembl IDs were used to map Atoh1 targetome genes to the pcHi-C dataset. Targetome Refseq IDs were mapped to protein coding Ensembl gene identifiers and mgi symbols using biomart (Ensembl 86). Where an Ensembl gene id or mgi symbol was not automatically detected, manual searching of the Refseq ID was performed using NCBI to identify an appropriate protein coding Ensembl gene ID or to assess if that Refseq ID was defunct. If more than one Ensembl ID was identified for a Refseq entry the protein-coding version of that Ensembl ID was selected.

#### Genome Visualisation

Coverage tracks and PCHiC interaction arcs were visualised using the Integrative Genomics Viewer web application (IGV-Web) (Robinson *et al*, 2011). Unless otherwise stated the transcript annotation displayed are from Ensembl 81 gtf file, that was filtered to only keep transcripts where the gencode tag= ‘basic’.

## Author contributions

KLHR performed the pcHi-C experiments under supervision of CSO, with capture probe library designed and provided by HY and BJPH. CG performed bioinformatic analysis under supervision of DS. DW and MUH performed the co-immunoprecipitation experiments. The study was conceived and designed by MAB, CSO and DS. The manuscript was written by KLHR, CG and MAB with input from all authors.

## Acknowledgements

We thank Chris Ponting and CGAT fellows who initiated the bioinformatic analysis of pcHi-C data and Daniel Fults (University of Utah) for the SHH-NPD cell line We acknowledge funding from the MRC (MR/K022377/1 and MR/Y008170/1 to MAB, G1000902 to DS and CG) and Wellcome Trust (224619/Z/21/Z to DEW).

## Data availability

Raw sequencing reads have been deposited to the Gene Expression Omnibus and will be made available upon publication. Details of code and pipelines used for analysis can be found at (https://github.com/sims-lab/GranuleProgenitor_pcHiC).

---

**Suppl Figure 1.**
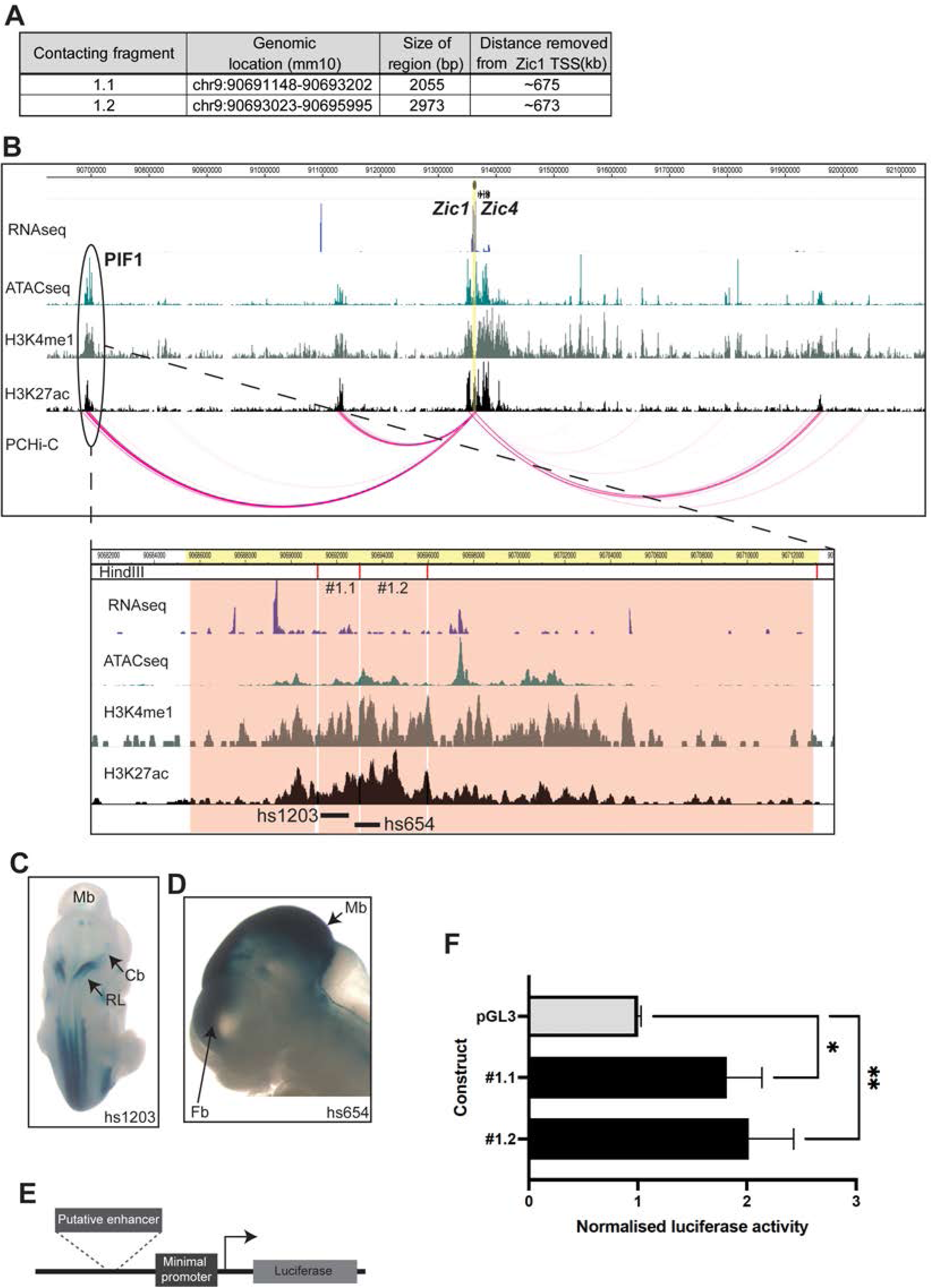
Putative enhancer elements within PIFs exhibit enhancer activity in GCps. **(A)** Table showing the genomoic location, size and difference from Zic1 TSS of two putative enhancer-containing fragments from PIF1 (Fig. 1G). **(B)** Diagram of *Zic1/4* locus with ATAC-seq, histone modification ChIP-seq and pcHi-C interactions shown in pink arcs. Note the location of PIF1, and the location of fragments #1, .1 and #1.2 in the magnified view below. Note the overlaps with VISTA enhancers hs1203 and hs654. **(C,D)** Enhancer activity of hindbrain VISTA enhancers in E11.5 mouse embryos. Mb=midbrain, Cb=cerebellum, RL-rhombic lip, Fb=forebrain. **(E)** Diagram of luciferase reporter construct. **(F)** Normalised luciferase activity of empty vector (pGL3) and enhancer fragments in SHH-NPD cells. *p<0.05, **p<0.01, t-test.

**Suppl Figure 2.**
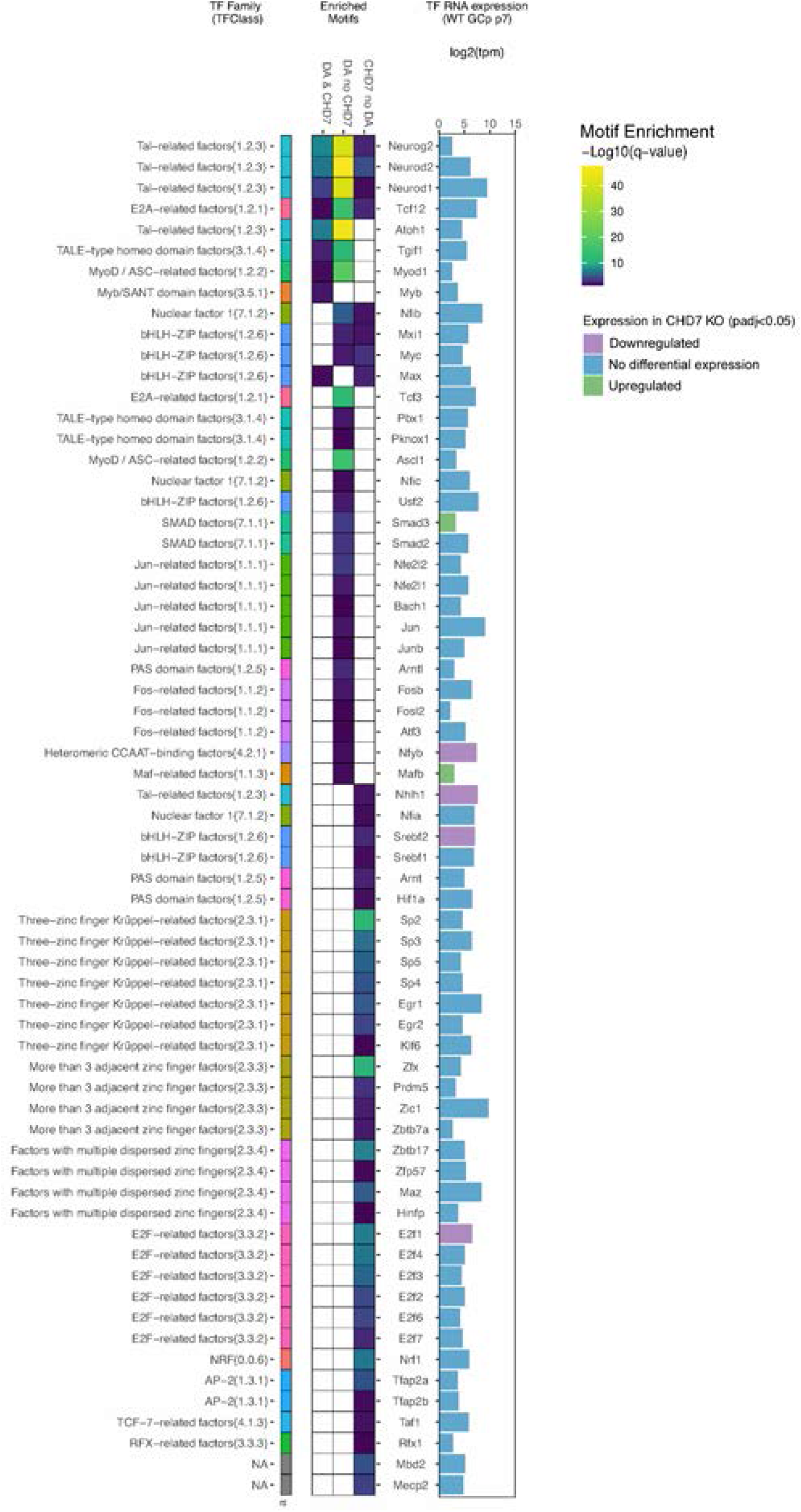
CHD7-regulated enhancers are enriched for proneural transcription factor motifs. A comparison of HOCOMOCO Mouse v11 CORE database transcription factor motifs enrichened within regulatory elements linked to CHD7 by SEA tool from MEME suite. Left panel describes TFClass family each transcription factor belongs to. Middle heatmap of q-values for significant SEA motif-enrichment results (P<0.05, Q<0.05) for genomic regions with ATAC peaks that are direct targets of CHD7 with CHD7 occupancy and change in accessibility in C*hd7*-deficient GCps (DA & CHD7), those that alter accessibility upon CHD7 depletion but do not display CHD7 occupancy in WT (DA no CHD7) and those which display CHD7 recruitment but do not alter accessibility upon CHD7 depletion (CHD7 no DA). White boxes denote no statistically significant enrichment. RNA expression level of transcription factors in WT GCps (n=2) is shown in right panel and color of bar indicates if the RNA expression level of that factor is unchanged in Chd7-depleted GCps (blue), upregulated (green) or downregulated (purple).

**Suppl Figure 3.**
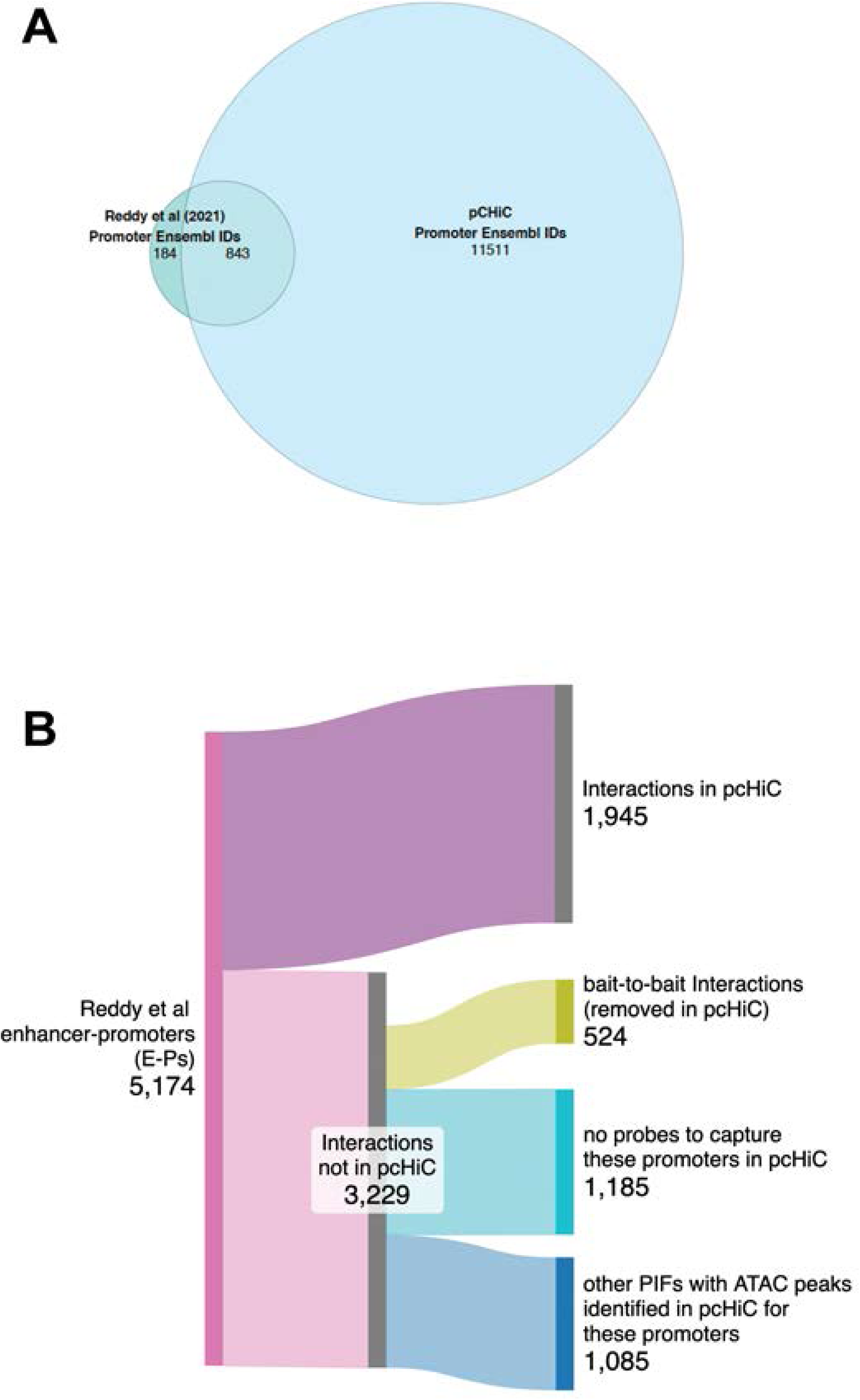
Comparison between our pcHi-C data from P7 mouse GCps and Hi-C data from P4 mouse cerebellum by Reddy et al. **(A)** Overlap of promoters with distal interactions identified in Reddy et al (green) vs. pCHiC (blue). **(B)** Sankey diagram demonstrating the number of E-P interactions from Reddy et al dataset replicated in the pCHiC data along with regions that are unique to the Reddy et al data, illustrating the proportion of the the E-P interactions that we would not expect to replicate in our data due to the technical design of the capture experiment. A substantial subset of Reddy E-P interactions were not found in our data due to technical differences (promoters not captured in our study), bioinformatic reasons (removal of bait-to-bait interactions. For some of the Reddy E-Ps, no interactions were identified in our data, or different potential enhancers (PIFs) were found, which might reflect differences in cell type and developmental timing.

## References

Allen G, Courchesne E (2003) Differential effects of developmental cerebellar abnormality on cognitive and motor functions in the cerebellum: an fMRI study of autism. Am J Psychiatry 160: 262–273

Amemiya HM, Kundaje A, Boyle AP (2019) The ENCODE Blacklist: Identification of Problematic Regions of the Genome. Sci Rep 9: 9354

Aruga J, Minowa O, Yaginuma H, Kuno J, Nagai T, Noda T, Mikoshiba K (1998) Mouse Zic1 is involved in cerebellar development. J Neurosci 18: 284–293

Bailey TL, Grant CE (2021a) SEA: Simple Enrichment Analysis of motifs. BioRxiv

Bailey TL, Grant CE (2021b) SEA: Simple enrichment of motifs. bioRxiv

Bajpai R, Chen DA, Rada-Iglesias A, Zhang J, Xiong Y, Helms J, Chang CP, Zhao Y, Swigut T, Wysocka J (2010) CHD7 cooperates with PBAF to control multipotent neural crest formation. Nature 463: 958–962

Basson MA, Wingate RJ (2013) Congenital hypoplasia of the cerebellum: developmental causes and behavioral consequences. Frontiers in neuroanatomy 7: 29

Ben-Arie N, Bellen HJ, Armstrong DL, McCall AE, Gordadze PR, Guo Q, Matzuk MM, Zoghbi HY (1997) Math1 is essential for genesis of cerebellar granule neurons. Nature 390: 169–172

Blank MC, Grinberg I, Aryee E, Laliberte C, Chizhikov VV, Henkelman RM, Millen KJ (2011) Multiple developmental programs are altered by loss of Zic1 and Zic4 to cause Dandy-Walker malformation cerebellar pathogenesis. Development 138: 1207–1216

Bolger AM, Lohse M, Usadel B (2014) Trimmomatic: a flexible trimmer for Illumina sequence data. Bioinformatics 30: 2114–2120

Bouwman BA, de Laat W (2015) Getting the genome in shape: the formation of loops, domains and compartments. Genome Biol 16: 154

Bray NL, Pimentel H, Melsted P, Pachter L (2016) Near-optimal probabilistic RNA-seq quantification. Nat Biotechnol 34: 525–527

Buenrostro JD, Wu B, Chang HY, Greenleaf WJ (2015) ATAC-seq: A Method for Assaying Chromatin Accessibility Genome-Wide. Current protocols in molecular biology / edited by Frederick M Ausubel [et al] 109: 21 29 21–29

Bulger M, Groudine M (2010) Enhancers: the abundance and function of regulatory sequences beyond promoters. Dev Biol 339: 250–257

Cairns J, Freire-Pritchett P, Wingett SW, Várnai C, Dimond A, Plagnol V, Zerbino D, Schoenfelder S, Javierre B-M, Osborne C et al (2016) CHiCAGO: robust detection of DNA looping interactions in Capture Hi-C data. Genome Biology 17: 127

Calo E, Wysocka J (2013) Modification of enhancer chromatin: what, how, and why? Mol Cell 49: 825–837

Chen Z, Snetkova V, Bower G, Jacinto S, Clock B, Dizehchi A, Barozzi I, Mannion BJ, Alcaina-Caro A, Lopez-Rios J et al (2024) Increased enhancer-promoter interactions during developmental enhancer activation in mammals. Nat Genet 56: 675–685

Consalez GG, Goldowitz D, Casoni F, Hawkes R (2020) Origins, Development, and Compartmentation of the Granule Cells of the Cerebellum. Front Neural Circuits 14: 611841

Corrales JD, Blaess S, Mahoney EM, Joyner AL (2006) The level of sonic hedgehog signaling regulates the complexity of cerebellar foliation. Development 133: 1811–1821

Corrales JD, Rocco GL, Blaess S, Guo Q, Joyner AL (2004) Spatial pattern of sonic hedgehog signaling through Gli genes during cerebellum development. Development 131: 5581–5590

Creyghton MP, Cheng AW, Welstead GG, Kooistra T, Carey BW, Steine EJ, Hanna J, Lodato MA, Frampton GM, Sharp PA et al (2010) Histone H3K27ac separates active from poised enhancers and predicts developmental state. Proc Natl Acad Sci U S A 107: 21931–21936

D’Mello AM, Stoodley CJ (2015) Cerebro-cerebellar circuits in autism spectrum disorder. Front Neurosci 9: 408

Dalby M, Rennie S, Andersson R ANTOM5 transcribed enhancers in mm10 (2018) (https://zenodo.org/records/1411211) [DATASET]

Danecek P, Bonfield JK, Liddle J, Marshall J, Ohan V, Pollard MO, Whitwham A, Keane T, McCarthy SA, Davies RM et al (2021) Twelve years of SAMtools and BCFtools. Gigascience 10

Dekker J, Rippe K, Dekker M, Kleckner N (2002) Capturing chromosome conformation. Science 295: 1306–1311

Dignam JD, Lebovitz RM, Roeder RG (1983) Accurate transcription initiation by RNA polymerase II in a soluble extract from isolated mammalian nuclei. Nucleic Acids Res 11: 1475–1489

Dobin A, Davis CA, Schlesinger F, Drenkow J, Zaleski C, Jha S, Batut P, Chaisson M, Gingeras TR (2013) STAR: ultrafast universal RNA-seq aligner. Bioinformatics 29: 15–21

Engelen E, Akinci U, Bryne JC, Hou J, Gontan C, Moen M, Szumska D, Kockx C, van Ijcken W, Dekkers DH et al (2011) Sox2 cooperates with Chd7 to regulate genes that are mutated in human syndromes. Nat Genet 43: 607–611

Fatemi SH, Aldinger KA, Ashwood P, Bauman ML, Blaha CD, Blatt GJ, Chauhan A, Chauhan V, Dager SR, Dickson PE et al (2012) Consensus Paper: Pathological Role of the Cerebellum in Autism. Cerebellum

Feng W, Kawauchi D, Korkel-Qu H, Deng H, Serger E, Sieber L, Lieberman JA, Jimeno-Gonzalez S, Lambo S, Hanna BS et al (2017) Chd7 is indispensable for mammalian brain development through activation of a neuronal differentiation programme. Nature communications 8: 14758

Freire-Pritchett P, Schoenfelder S, Varnai C, Wingett SW, Cairns J, Collier AJ, Garcia-Vilchez R, Furlan-Magaril M, Osborne CS, Fraser P et al (2017) Global reorganisation of cis-regulatory units upon lineage commitment of human embryonic stem cells. eLife 6

Gates LA, Foulds CE, O’Malley BW (2017) Histone Marks in the ’Driver’s Seat’: Functional Roles in Steering the Transcription Cycle. Trends Biochem Sci 42: 977–989

Grinberg I, Northrup H, Ardinger H, Prasad C, Dobyns WB, Millen KJ (2004) Heterozygous deletion of the linked genes ZIC1 and ZIC4 is involved in Dandy-Walker malformation. Nat Genet 36: 1053–1055

Hatten ME (1985) Neuronal regulation of astroglial morphology and proliferation in vitro. J Cell Biol 100: 384–396

Heinz S, Benner C, Spann N, Bertolino E, Lin YC, Laslo P, Cheng JX, Murre C, Singh H, Glass CK (2010) Simple combinations of lineage-determining transcription factors prime cis-regulatory elements required for macrophage and B cell identities. Mol Cell 38: 576–589

Hinrichs AS, Karolchik D, Baertsch R, Barber GP, Bejerano G, Clawson H, Diekhans M, Furey TS, Harte RA, Hsu F et al (2006) The UCSC Genome Browser Database: update 2006. Nucleic Acids Res 34: D590–598

Hong SE, Shugart YY, Huang DT, Shahwan SA, Grant PE, Hourihane JO, Martin ND, Walsh CA (2000) Autosomal recessive lissencephaly with cerebellar hypoplasia is associated with human RELN mutations. Nat Genet 26: 93–96

Hsu J, Huang HT, Lee CT, Choudhuri A, Wilson NK, Abraham BJ, Moignard V, Kucinski I, Yu S, Hyde RK et al (2020) CHD7 and Runx1 interaction provides a braking mechanism for hematopoietic differentiation. Proc Natl Acad Sci U S A 117: 23626–23635

Ignatiadis N, Klaus B, Zaugg JB, Huber W (2016) Data-driven hypothesis weighting increases detection power in genome-scale multiple testing. Nat Methods 13: 577–580

Kadauke S, Blobel GA (2009) Chromatin loops in gene regulation. Biochim Biophys Acta 1789: 17–25

Kelly E, Meng F, Fujita H, Morgado F, Kazemi Y, Rice LC, Ren C, Escamilla CO, Gibson JM, Sajadi S et al (2020) Regulation of autism-relevant behaviors by cerebellar-prefrontal cortical circuits. Nat Neurosci 23: 1102–1110

Kent WJ, Zweig AS, Barber G, Hinrichs AS, Karolchik D (2010) BigWig and BigBed: enabling browsing of large distributed datasets. Bioinformatics 26: 2204–2207

Klisch TJ, Xi Y, Flora A, Wang L, Li W, Zoghbi HY (2011) In vivo Atoh1 targetome reveals how a proneural transcription factor regulates cerebellar development. Proc Natl Acad Sci U S A 108: 3288–3293

Kolberg L, Raudvere U, Kuzmin I, Adler P, Vilo J, Peterson H (2023) g:Profiler-interoperable web service for functional enrichment analysis and gene identifier mapping (2023 update). Nucleic Acids Res 51: W207–W212

Krizhanovsky V, Soreq L, Kliminski V, Ben-Arie N (2006) Math1 target genes are enriched with evolutionarily conserved clustered E-box binding sites. J Mol Neurosci 28: 211–229

Langmead B, Salzberg SL (2012) Fast gapped-read alignment with Bowtie 2. Nat Methods 9: 357-359

Langmead B, Trapnell C, Pop M, Salzberg SL (2009) Ultrafast and memory-efficient alignment of short DNA sequences to the human genome. Genome Biol 10: R25

Li G, Ruan X, Auerbach RK, Sandhu KS, Zheng M, Wang P, Poh HM, Goh Y, Lim J, Zhang J et al (2012) Extensive promoter-centered chromatin interactions provide a topological basis for transcription regulation. Cell 148: 84–98

Lieberman-Aiden E, van Berkum NL, Williams L, Imakaev M, Ragoczy T, Telling A, Amit I, Lajoie BR, Sabo PJ, Dorschner MO et al (2009) Comprehensive mapping of long-range interactions reveals folding principles of the human genome. Science 326: 289–293

Love MI, Huber W, Anders S (2014) Moderated estimation of fold change and dispersion for RNA-seq data with DESeq2. Genome Biol 15: 550

Machold R, Fishell G (2005) Math1 is expressed in temporally discrete pools of cerebellar rhombic-lip neural progenitors. Neuron 48: 17–24

Marsh APL, Edwards TJ, Galea C, Cooper HM, Engle EC, Jamuar SS, Meneret A, Moutard ML, Nava C, Rastetter A et al (2018) DCC mutation update: Congenital mirror movements, isolated agenesis of the corpus callosum, and developmental split brain syndrome. Hum Mutat 39: 23–39

Mifsud B, Tavares-Cadete F, Young AN, Sugar R, Schoenfelder S, Ferreira L, Wingett SW, Andrews S, Grey W, Ewels PA et al (2015) Mapping long-range promoter contacts in human cells with high-resolution capture Hi-C. Nat Genet 47: 598–606

Miyata T, Maeda T, Lee JE (1999) NeuroD is required for differentiation of the granule cells in the cerebellum and hippocampus. Genes Dev 13: 1647–1652

Pieper A, Rudolph S, Wieser GL, Gotze T, Miessner H, Yonemasu T, Yan K, Tzvetanova I, Castillo BD, Bode U et al (2019) NeuroD2 controls inhibitory circuit formation in the molecular layer of the cerebellum. Sci Rep 9: 1448

Quinlan AR, Hall IM (2010) BEDTools: a flexible suite of utilities for comparing genomic features. Bioinformatics 26: 841–842

Rad A, Altunoglu U, Miller R, Maroofian R, James KN, Caglayan AO, Najafi M, Stanley V, Boustany RM, Yesil G et al (2019) MAB21L1 loss of function causes a syndromic neurodevelopmental disorder with distinctive cerebellar, ocular, craniofacial and genital features (COFG syndrome). J Med Genet 56: 332–339

Ramirez F, Ryan DP, Gruning B, Bhardwaj V, Kilpert F, Richter AS, Heyne S, Dundar F, Manke T (2016) deepTools2: a next generation web server for deep-sequencing data analysis. Nucleic Acids Res 44: W160–165

Reddy NC, Majidi SP, Kong L, Nemera M, Ferguson CJ, Moore M, Goncalves TM, Liu HK, Fitzpatrick JAJ, Zhao G et al (2021) CHARGE syndrome protein CHD7 regulates epigenomic activation of enhancers in granule cell precursors and gyrification of the cerebellum. Nature communications 12: 5702

Robinson JT, Thorvaldsdottir H, Winckler W, Guttman M, Lander ES, Getz G, Mesirov JP (2011) Integrative genomics viewer. Nat Biotechnol 29: 24–26

Sanyal A, Lajoie BR, Jain G, Dekker J (2012) The long-range interaction landscape of gene promoters. Nature 489: 109–113

Schnetz MP, Handoko L, Akhtar-Zaidi B, Bartels CF, Pereira CF, Fisher AG, Adams DJ, Flicek P, Crawford GE, Laframboise T et al (2010) CHD7 targets active gene enhancer elements to modulate ES cell-specific gene expression. PLoS Genet 6: e1001023

Schoenfelder S, Fraser P (2019) Long-range enhancer-promoter contacts in gene expression control. Nat Rev Genet 20: 437–455

Schoenfelder S, Furlan-Magaril M, Mifsud B, Tavares-Cadete F, Sugar R, Javierre BM, Nagano T, Katsman Y, Sakthidevi M, Wingett SW et al (2015) The pluripotent regulatory circuitry connecting promoters to their long-range interacting elements. Genome Res 25: 582–597

Siersbaek R, Madsen JGS, Javierre BM, Nielsen R, Bagge EK, Cairns J, Wingett SW, Traynor S, Spivakov M, Fraser P et al (2017) Dynamic Rewiring of Promoter-Anchored Chromatin Loops during Adipocyte Differentiation. Mol Cell 66: 420–435 e425

Soneson C, Love MI, Robinson MD (2015) Differential analyses for RNA-seq: transcript-level estimates improve gene-level inferences. F1000Res 4: 1521

Stathopoulou A, Wang P, Thellier C, Kelly RG, Zheng D, Scambler PJ (2023) CHARGE syndrome-associated CHD7 acts at ISL1-regulated enhancers to modulate second heart field gene expression. Cardiovasc Res

Swanson DJ, Goldowitz D (2011) Experimental Sey mouse chimeras reveal the developmental deficiencies of Pax6-null granule cells in the postnatal cerebellum. Dev Biol 351: 1–12

Villiers W, Kelly A, He X, Kaufman-Cook J, Elbasir A, Bensmail H, Lavender P, Dillon R, Mifsud B, Osborne CS (2023) Multi-omics and machine learning reveal context-specific gene regulatory activities of PML::RARA in acute promyelocytic leukemia. Nature communications 14: 724

Visel A, Minovitsky S, Dubchak I, Pennacchio LA (2007) VISTA Enhancer Browser--a database of tissue-specific human enhancers. Nucleic Acids Res 35: D88–92

Wang SS, Kloth AD, Badura A (2014) The cerebellum, sensitive periods, and autism. Neuron 83: 518–532

Whittaker DE, Kasah S, Donovan APA, Ellegood J, Riegman KLH, Volk HA, McGonnell I, Lerch JP, Basson MA (2017a) Distinct cerebellar foliation anomalies in a CHD7 haploinsufficient mouse model of CHARGE syndrome. American Journal of Medical Genetics Part C: Seminars in Medical Genetics 175

Whittaker DE, Riegman KL, Kasah S, Mohan C, Yu T, Pijuan-Sala B, Hebaishi H, Caruso A, Marques AC, Michetti C et al (2017b) The chromatin remodeling factor CHD7 controls cerebellar development by regulating reelin expression. J Clin Invest 127: 874–887

Wingett S, Ewels P, Furlan-Magaril M, Nagano T, Schoenfelder S, Fraser P, Andrews S (2015) HiCUP: pipeline for mapping and processing Hi-C data. F1000Res 4: 1310–1310

Yamada M, Seto Y, Taya S, Owa T, Inoue YU, Inoue T, Kawaguchi Y, Nabeshima Y, Hoshino M (2014) Specification of spatial identities of cerebellar neuron progenitors by ptf1a and atoh1 for proper production of GABAergic and glutamatergic neurons. J Neurosci 34: 4786–4800

Yu G, Wang LG, He QY (2015) ChIPseeker: an R/Bioconductor package for ChIP peak annotation, comparison and visualization. Bioinformatics 31: 2382–2383

Yu T, Meiners LC, Danielsen K, Wong MT, Bowler T, Reinberg D, Scambler PJ, van Ravenswaaij-Arts CM, Basson MA (2013) Deregulated FGF and homeotic gene expression underlies cerebellar vermis hypoplasia in CHARGE syndrome. eLife 2: e01305

Yun H, Narayan N, Vohra S, Giotopoulos G, Mupo A, Madrigal P, Sasca D, Lara-Astiaso D, Horton SJ, Agrawal-Singh S et al (2021a) Mutational synergy during leukemia induction remodels chromatin accessibility, histone modifications and three-dimensional DNA topology to alter gene expression. Nat Genet 53: 1443–1455

Yun H, Narayan N, Vohra S, Giotopoulos G, Mupo A, Madrigal P, Sasca D, Lara-Astiaso D, Horton SJ, Agrawal-Singh S et al (2021b) Mutational synergy during leukemia induction remodels chromatin accessibility, histone modifications and three-dimensional DNA topology to alter gene expression. Nature Genetics 53: 1443–1455

Zerbino DR, Johnson N, Juetteman T, Sheppard D, Wilder SP, Lavidas I, Nuhn M, Perry E, Raffaillac-Desfosses Q, Sobral D et al (2016) Ensembl regulation resources. Database (Oxford) 2016

Zerbino DR, Johnson N, Juettemann T, Wilder SP, Flicek P (2014) WiggleTools: parallel processing of large collections of genome-wide datasets for visualization and statistical analysis. Bioinformatics 30: 1008–1009

Zhang Y, Liu T, Meyer CA, Eeckhoute J, Johnson DS, Bernstein BE, Nusbaum C, Myers RM, Brown M, Li W et al (2008) Model-based analysis of ChIP-Seq (MACS). Genome Biol 9: R137

